# Pyrogallol Modulates Abscopal Tumour and Gut Microbial Responses to Localized Irradiation in an Ehrlich Ascites Carcinoma Model

**DOI:** 10.64898/2026.08.16.745132

**Authors:** Sayantanee Ray, Rubin Nishanth Armstrong, Devipriya Nagarajan, Prakash Shankaran

## Abstract

Radiotherapy’s clinical utility is often limited by radio-resistance, enterotoxicity, and intestinal dysbiosis. This study evaluated pyrogallol—a plant-derived vicinal trihydroxybenzene—as a dual-action radiosensitizer and mucosal protectant in an Ehrlich ascites carcinoma (EAC) BALB/c mouse model subjected to targeted LINAC irradiation (8 Gy). By combining transcriptomic profiling with whole-genome metagenomic sequencing, we interrogated the underlying host-microbiome interactions. Pyrogallol co-treatment significantly augmented radiotherapeutic efficacy, driving marked tumour regression through the upregulation of pro-apoptotic effectors (*Bax*, *Casp3*, *Casp7*) and *p53*-mediated tumour suppressors (*Tp53*, *p21*), alongside *Bcl2* repression. Concurrently, pyrogallol blunted oncogenic progression by arresting proliferation (*Cdk4*, *Pcna*), inhibiting epithelial-mesenchymal transition (*N-cadherin*, *vimentin*), downregulating fibrotic remodelling (*Tgf-β*, *Col1A1*, *Fibronectin*), and attenuating radiation-induced pro-inflammatory cytokine surges (*Il-1α*, *Il-6*, *Il-12*). At the gut interface, radiation degraded colonization resistance by depleting homeostatic short-chain fatty acid producers and *Clostridium scindens*, while fuelling pathobiont blooms (*Acinetobacter baumannii*, *Clostridioides difficile*). Pyrogallol reversed this dysbiosis through a distinct ecological shift; despite a reduction in total species richness, the intestinal niche became dominated by the next-generation probiotic *Parabacteroides distasonis* (∼94% relative abundance; Berger-Parker index: 0.94). Integrated Spearman’s rank correlations demonstrated that host proliferative, EMT, fibrotic, and inflammatory markers aligned positively with pathobiont clusters (*Bacteroides caecimuris*, *B. faecium*, *A. baumannii*). Conversely, tumour regression and anti-inflammatory signatures correlated strongly with pathobiont restriction and *P. distasonis* enrichment. Overall, pyrogallol emerges as a compelling therapeutic adjuvant that synergistically enhances tumour radiosensitivity while remodelling the gut microbiome into a protective, anti-inflammatory state.

## 1. Introduction

Cancer remains one of the leading causes of morbidity and mortality worldwide, driven by unchecked cellular proliferation, genomic instability, and altered survival signalling (1,2). Among conventional treatment modalities, radiotherapy plays a central role in managing localized and metastatic solid tumours by inducing direct DNA double-strand breaks and indirect reactive oxygen species (ROS)-mediated genomic damage (3,4). However, two major hurdles limit its clinical success: acquired radio-resistance within the tumour microenvironment mediated by enhanced DNA repair, hypoxia, and epithelial-mesenchymal transition (EMT) (5) and collateral damage to healthy tissues (6).

When directed at abdominal or pelvic sites, ionizing radiation severely injures the rapidly proliferating intestinal epithelium. This gastrointestinal toxicity triggers localized inflammation, tissue fibrosis, and EMT-driven remodelling. Crucially, radiation disrupts the bidirectional host microbiome axis, causing profound gut bacteriome restructuring (7,8). This dysbiosis depletes beneficial, short-chain fatty acid (SCFA) producing commensals and drives the expansion of opportunistic pathogens, inducing a systemic pro-inflammatory state (9,10).

To overcome radio-resistance while mitigating normal tissue injury, natural polyphenols have gained significant attention as radiosensitizers and radioprotectors (11). Pyrogallol (1,2,3-trihydroxybenzene), a plant-derived vicinal trihydroxybenzene, possesses potent redox-modulating, anti-inflammatory, and selective antitumor properties. While pyrogallol is known to alter intracellular ROS dynamics and trigger apoptosis in malignant cells (12), its systemic capacity to simultaneously act as an oncological radiosensitizer and a protective agent against radiation-induced gut injury remains unexplored.

Understanding these multi-organ dynamics requires resolving host molecular shifts alongside species-level microbial alterations. Traditional short-read sequencing often lacks the resolution required to identify specific commensal or pathogenic strains. In contrast, third-generation Oxford Nanopore Technology (ONT) long-read metagenomics enables assembly-free, species-level profiling. To economically navigate animal cohort variability prior to large-scale trials, initial baseline survey using pooled metagenomic workflow offer an exploratory strategic approach.

To bridge these knowledge gaps, this study evaluates the radio-sensitizing profile of pyrogallol in a subcutaneous Ehrlich Ascites Carcinoma (EAC) BALB/c mouse model. Notably, localized thoracic X-ray irradiation was chosen as it triggers systemic immune and an abscopal effect (13) that can regress distant subcutaneous Ehrlich ascites carcinoma (EAC) tumours. Concurrently, this exposure induces out-of-field gut dysbiosis (14) and the mucosal barrier compromise mediated by circulating pro-inflammatory cytokine cascades (15), making it an ideal model to study both tumor response and intestinal injury simultaneously. Using quantitative real-time PCR (qPCR), we map small intestinal gene expression across apoptotic (*Bcl2, Bax, Caspases*), cell-cycle (*Cdk4, p21, Tp53*), proliferative (*Pcna*), fibrotic (*Tgf-β, Col1A1, Fibronectin*), and inflammatory (*Il-1α, Il-6, Il-12*) pathways. Concurrently, we perform exploratory ONT MinION metagenomic profiling on pooled faecal samples to map species-level taxonomic shifts. Through integrated Spearman’s correlation analysis, we establish key baseline relationships between host tissue markers and emerging gut microbiome clusters, providing a foundational framework for pyrogallol as a dual-action therapeutic adjuvant.

## 2. Materials and methods

### 2.1. Animal model and cancer induction

Male BALB/c mice aged 5 weeks were obtained from Central Animal Facility at SASTRA Deemed University, Tamil Nadu, India. The animals were then acclimatized for 7 days under controlled conditions with 12 h dark/light cycle. All experimental procedures conducted were approved by the Institutional Animal Ethics Committee (IAEC), Sastra Deemed University, Tamil Nadu, India (Approval No. 718/SASTRA/IAEC/RPP). A total of 30 mice were divided into five groups of six animals each. Group I: Control (n=6), Group II: Cancer (n=6), Group III: cancer + radiation 8 Gy (C+R) (n=6), Group IV: cancer + radiation 8 Gy + pyrogallol (20 mg/kg) (C+R+Py) (n=6), Group V: cancer + pyrogallol (20 mg/kg) (C+Py) (n=6). Cancer was induced in Group II mice by subcutaneously injecting EAC cells of density 1 x 10^6^ and observed for 7 days for tumour development. Group IV animals were intraperitoneally administered with pyrogallol (20 mg/kg) for 7 consecutive days before X-ray irradiation. At the end of experimental period (23 days), mice were euthanized using CO_2_ induction method. Small intestine was subsequently isolated, stored at -80 °C, and used for further experiments.

### 2.2. Radiation exposure

Mice were anaesthetized with ketamine (30 mg/kg), placed supine on a foam holder with the thoracic region immobilized using plastic bandages, and exposed to a single fraction of 8 Gy localized X-ray irradiation using a Linear Accelerator (LINAC) at Vishnu Cancer Centre, Thanjavur, India.

### 2.3. RNA isolation and cDNA synthesis

Total RNA was extracted from small intestine (250 mg, n=3 Biological triplicate; samples have been collected from 3 mice out of 6 mice) using TRIzol RNAiso Plus (#9108) (TaKaRa, Shiga, Japan). The purity and concentration of RNA was determined using nanodrop spectrophotometer (Thermo Scientific NanoDrop 2000/2000c spectrophotometer). A total of 1 µg of RNA was converted to cDNA using PrimeScript RT reagent kit (#RR037A) (TaKaRa, Shiga, Japan). The reaction was set up at 37 °C for 10 min, followed by 81 °C for 5 s to deactivate the enzyme in RT-PCR system (Bio-Rad CFX Opus 96, USA)

### 2.4. Real-time PCR

Real-time PCR was carried out using TB Green Premix Ex Taq II (Tli RNase H Plus) (#RR820A, TaKaRa) in Bio-Rad CFX Opus 96 system. The q-PCR reaction was set up at 95°C for 3 minutes and 15 seconds, followed by melting temperature (T_m_) for 1 minute for 39 cycles, after which 65°C was maintained for 5 seconds and 95°C for 50 seconds. The expressed gene products were normalized using β-actin as an internal control. The list of primers and their respective sequences were given in Table.

### 2.5. Statistical analysis

All experiments were performed in triplicates (n=3 independent experiments), results were expressed as mean ± standard deviation. One way ANOVA was used to analyse significant difference between multiple groups in GraphPad Prism 8.0. A p-value ≤ 0.05 was defined as statistically significant with significance levels mentioned as *p < 0.05, **p < 0.01, ***p < 0.001, and ****p < 0.0001.

### 2.6. DNA isolation and quantification

Following previously published studies (16) to economically evaluate representative baseline microbial shifts across the six experimental cohorts while intentionally averaging across individual biological variance to identify cohort-level trends, equal mass amounts of faecal pellets from 6 mice per group were pooled into composite cohort prior to total DNA extraction. On the final day of the study period (day 23) the pooled faecal samples were collected from each group and stored in -20°C freezer until further processing. Total DNA was extracted from faecal sample (200-250mg) using QIAmp Fast DNA Stool Mini Kit (Cat No 51604) (QIAGEN, Hilden, Germany) according to manufacturer protocol. The purity of DNA was determined using nanodrop spectrophotometer (Thermo Scientific NanoDrop 2000/2000c spectrophotometer) measuring the ratio of 260/280 and 260/230. Qubit dsDNA Quantification Assay kit (Cat No Q32851) (Invitrogen, Thermo Fischer Scientific) was used to achieve accurate and precise quantification of double stranded DNA from the samples; which were used for downstream processes.

### 2.7. Whole genome metagenome sequencing

Whole genome metagenome sequencing was performed using MinION Mk1B machine using MinION Flow cells (R10.4.1) and Native barcoding kit (SQK NBD114.24) (V 14 chemistry) from Oxford Nanopore Technology (ONT), United Kingdom. Library preparation including DNA repair and end preparation, barcoding and ligation were carried out according to ONT protocol using the enzymes from New England Biolabs. Raw data was generated by MinKNOW (Version 26.01.15) software as POD5 files and base calling was performed using high accuracy model of DORADO basecaller.

### 2.8. Data Analysis

Basecalled FASTQ files were analysed using EPI2ME’s assembly-free metagenomics workflow; which is a standalone desktop application provided by ONT. Taxonomic classification was performed with Kraken 2 using the Standard-8 reference database which comprises NCBI Refseq archaea, bacteria, viral, plasmid, human.

### 2.9. Diversity index calculation

#### 2.9.1. Alpha diversity

The Berger-Parker index, Chao1, Shannon diversity and Simpson index were calculated using R (Vegan) and plotted using ggplot2. Some sample groups exhibited lower total abundance, to account for uneven sequencing depth, all samples were rarefied to a depth of reads using the R package vegan (rarefy function).

#### 2.9.2. Beta diversity

Principal Coordinate Analysis (PCoA) plot using the Bray-Curtis dissimilarity formula visualizes multidimensional microbiome data into a 2D scatter plot; here it was calculated using R (Vegan) and plotted using ggplot2.

### 2.10. Spearman’s correlation calculation

Spearman’s correlation coefficient was calculated using Vegan package of R and heatmap was plotted using ComplexHeatmap. Normalized Z-score was used for each gene expression values and relative abundance of bacterial species.

## 3. Results

### 3.1. Pyrogallol induced apoptosis and sensitized the cancer-induced Balb/c mice to radiotherapy

The expression levels of anti-apoptotic gene *Bcl2* and pro-apoptotic genes including Bax, *caspase 3* and *caspase 7* were analyzed by RT-PCR to evaluate the modifications of apoptosis in different experimental groups. Our results indicated a significant upregulation of *Bcl2* in the cancer induced mice when compared to the control group, demonstrating enhanced anti-apoptotic signaling. Upon exposure to radiotherapy, a noticeable reduction in *Bcl2* expression was observed in the cancer induced mice suggesting that radiation treatment suppressed anti-apoptotic gene. Treatment with pyrogallol in combination with radiotherapy resulted in a further decrease in *Bcl2* gene expression, highlighting radio sensitizing potential of pyrogallol by mitigating anti-apoptotic mechanism. (Fig 1a)

**Figure 1a:**
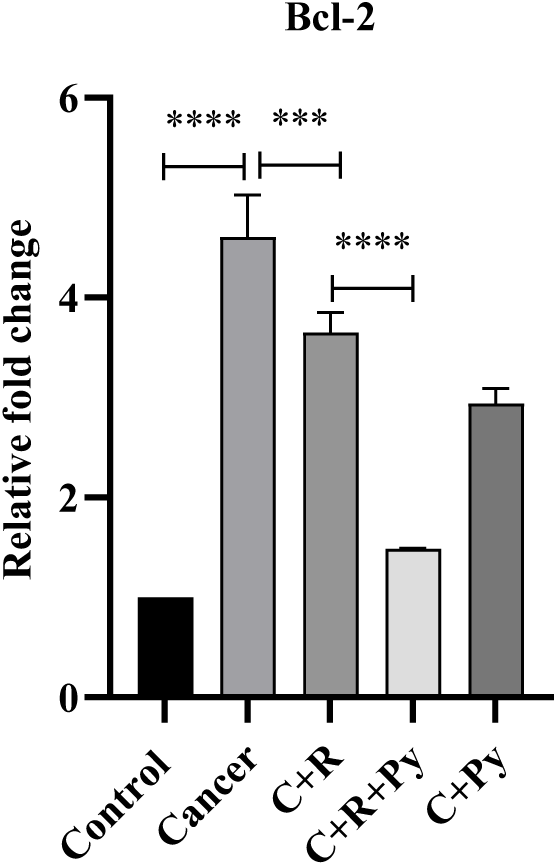
Relative expression profile of host *Bcl2* across experimental cohorts. (Real-time quantitative PCR analysis showing the relative fold change in *Bcl2* mRNA expression levels normalized to host reference gene across five experimental groups (n=3 from each group): Control, Cancer (EAC tumour), C+R (Cancer + Radiotherapy), C+R+Py (Cancer + Radiotherapy + Pyrogallol), and C+Py (Cancer + Pyrogallol). Data are presented as mean ± SD; Statistical significance was determined using one-way ANOVA *p < 0.05, **p < 0.01, ***p < 0.001, ****p < 0.0001; ns, not significant).

**Fig 1b, c, d:**
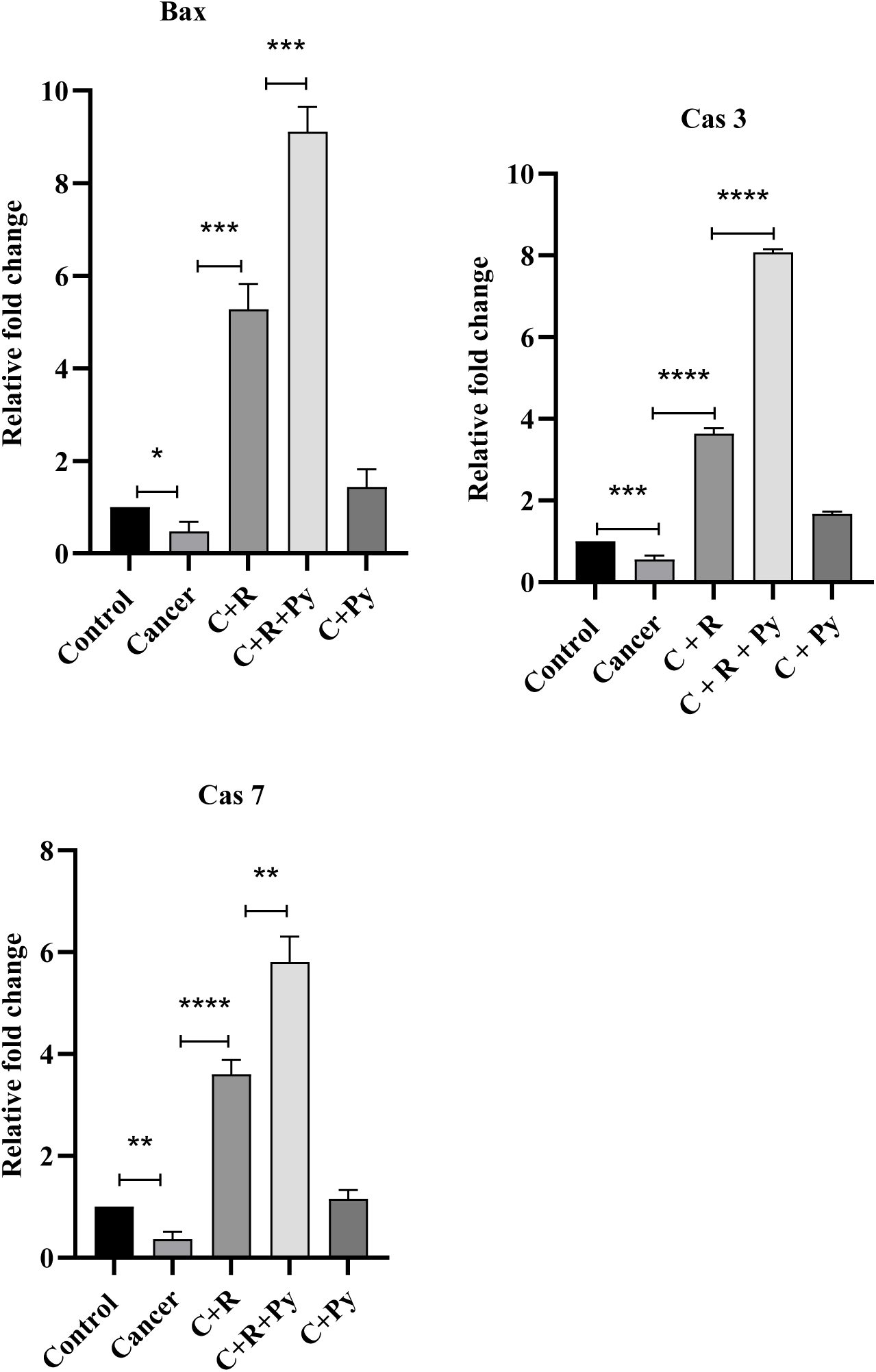
Relative expression profile of host *Bax, Cas3* and *Cas7* respectively across experimental cohorts. (Real-time quantitative PCR analysis showing the relative fold change in *Bax, Cas3* and *Cas7* mRNA expression levels normalized to host reference gene across five experimental groups (n=3 from each group): Control, Cancer (EAC tumour), C+R (Cancer + Radiotherapy), C+R+Py (Cancer + Radiotherapy + Pyrogallol), and C+Py (Cancer + Pyrogallol). Data are presented as mean ± SD; Statistical significance was determined using one-way ANOVA *p < 0.05, **p < 0.01, ***p < 0.001, ****p < 0.0001; ns, not significant).

In contrast, the expression level of pro-apoptotic genes *Bax*, *caspase-3* and *caspase-7* were significantly downregulated in the cancer-induced mice compared with the control group. Radiotherapy treatment noticeable increased the expression of these apoptotic genes shown in Figure 1b, c, d. Co-treatment with radiation and pyrogallol, upregulated the apoptotic gene expression indicating the radio-sensitization activity of pyrogallol. Animals treated with pyrogallol only did not exhibit any significant alterations in the expression of apoptotic genes.

### 3.2. Pyrogallol-mediated Cdk4 suppression promotes radiotherapy sensitivity in cancer-induced Balb/c mice

The *Cdk4* (cyclin-dependent kinase 4) gene is essential for regulating the cell cycle. *Cdk4* pairs with Cyclin D to drive cells through the G_1 phase (preparation for DNA replication) into the S phase (DNA synthesis). It achieves this by phosphorylating the retinoblastoma (RB) protein, which releases E2F transcription factors and allows the cell to divide. Upregulation of *Cdk4* gene was observed in cancer-induced Balb/c mice compared to the control group indicating enhanced cell cycle progression associated with tumor development. Treatment with radiotherapy resulted in significant reduction in *Cdk4* gene expression compared with the untreated cancer group indicating that radiotherapy suppressed *Cdk4* mediated cell cycle progression as shown in Figure 2a. Notably, the combined treatment of radiotherapy and pyrogallol further decreased *Cdk4* expression compared with the radiotherapy-only group showing an inhibitory effect on *Cdk4*-mediated cell cycle regulation. Mice treated with pyrogallol alone showed similar levels of *Cdk4* to control group.

**Figure 2a:**
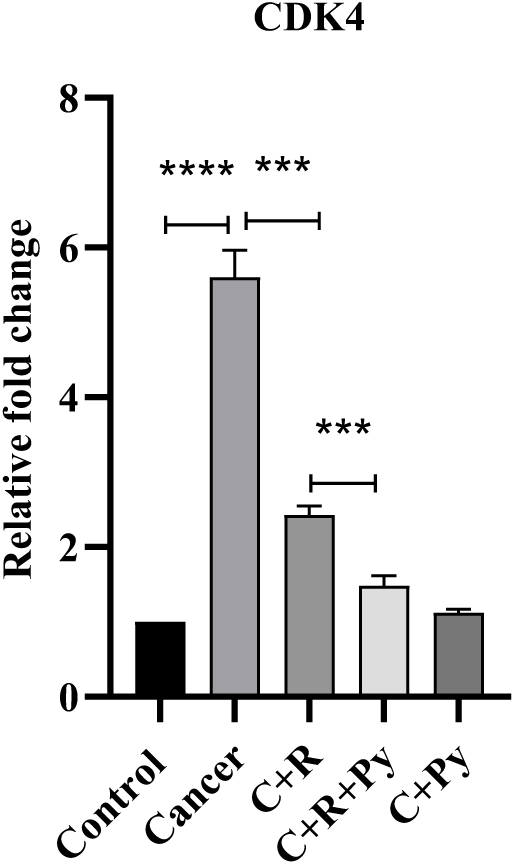
Relative expression profile of host *Cdk4* across experimental cohorts. (Real-time quantitative PCR analysis showing the relative fold change in *Cdk4* mRNA expression levels normalized to host reference gene across five experimental groups (n=3 from each group): Control, Cancer (EAC tumour), C+R (Cancer + Radiotherapy), C+R+Py (Cancer + Radiotherapy + Pyrogallol), and C+Py (Cancer + Pyrogallol). Data are presented as mean ± SD; Statistical significance was determined using one-way ANOVA *p < 0.05, **p < 0.01, ***p < 0.001, ****p < 0.0001; ns, not significant).

### 3.3. Pyrogallol modulates Pcna gene expression and reduced tumor cell proliferation in cancer-induced Balb/c mice

The *Pcna* (Proliferating Cell Nuclear Antigen) gene provides instructions for making the *Pcna* protein. This protein acts as an essential cofactor (processivity factor) for DNA polymerases during cell division, ensuring the DNA is copied quickly and accurately. It also recruits specialized proteins to fix damaged or mutated DNA and interacts with checkpoint proteins to help the cell decide whether to divide, pause for repair, or undergo cell death. The relative gene expression of *Pcna* was evaluated to determine the effect of radiotherapy and pyrogallol treatment on tumor-cell proliferation in cancer-induced Balb/c mice. Our results showed significant increase in *Pcna* gene expression in cancer-induced group compared with the control group indicating enhanced cancer cell proliferation. Following radiotherapy treatment, *Pcna* expression was reduced (Figure: 2(B)) compared with the untreated cancer-induced group suggesting that radiotherapy suppressed tumor proliferation. A further reduction in *Pcna* expression was observed in the cancer-induced mice group treated with a combination of radiotherapy and pyrogallol. Mice treated with pyrogallol alone showed a moderate reduction in *Pcna* expression compared with the cancer-induced group. However, *Pcna* expression was similar in drug only group and the control group.

**Fig 2b:**
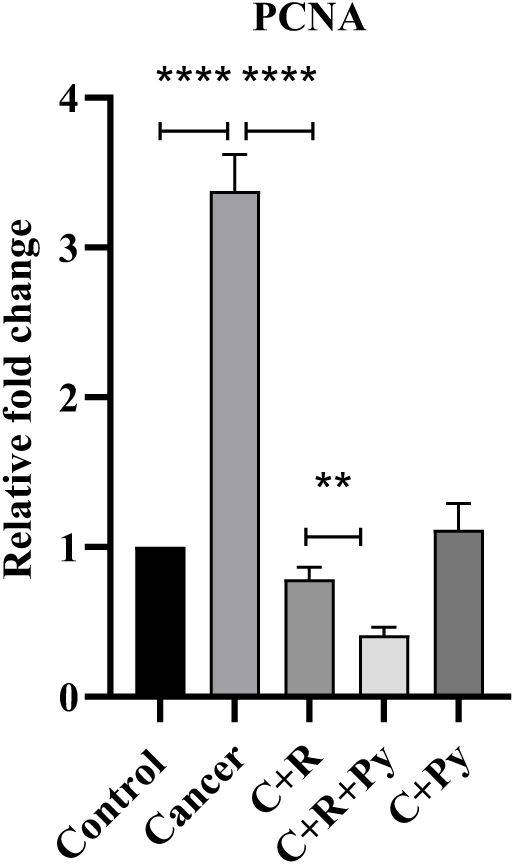
Relative expression profile of host *Pcna* across experimental cohorts. (Real-time quantitative PCR analysis showing the relative fold change in *Pcna* mRNA expression levels normalized to host reference gene across five experimental groups (n=3 from each group): Control, Cancer (EAC tumour), C+R (Cancer + Radiotherapy), C+R+Py (Cancer + Radiotherapy + Pyrogallol), and C+Py (Cancer + Pyrogallol). Data are presented as mean ± SD; Statistical significance was determined using one-way ANOVA *p < 0.05, **p < 0.01, ***p < 0.001, ****p < 0.0001; ns, not significant).

### 3.4. Pyrogallol reduced fibrotic gene expression in cancer-induced Balb/c mice during radiotherapy

*Tgf-β* (Transforming growth factor β) is a master regulatory protein and signalling molecule. It directs cell growth, tissue repair, and immune regulation. *Col1A1* (Collagen Type I Alpha 1) provides instructions for making part of Type I collagen, the most abundant protein in the body. *Fibronectin* is a glycoprotein that acts as biological glue. It binds to cells and other extracellular matrix (ECM) proteins (like collagen) to help them stick together. The relative gene expressions of *Tgf-β*, *Col1A1* and *Fibronectin*, key regulators of fibrosis and tissue remodeling, was analyzed and our results demonstrated a significant increase in *Tgf-β* expression in the cancer-induced group when compared to control group, indicating increased fibrosis signaling associated with tumor progression. Radiotherapy treated cancer-induced mice showed a notable reduction in all these fibrotic gene expressions compared to untreated cancer-induced group. However, the expression level remained higher than the control group. More reduction in *Tgf-β*, *Col1A1* and *Fibronectin* was observed in the combined treatment group that received radiotherapy and pyrogallol in Figure: 3a, b, c, highlighting that pyrogallol enhanced the suppressive effect of radiotherapy on fibrotic genes. Mice treated with pyrogallol alone showed a substantial reduction in *Tgf-β* expression but gene expression levels of *Fibronectin* and *Col1A1* were increased little bit compared to control group.

**Fig 3a, b, c:**
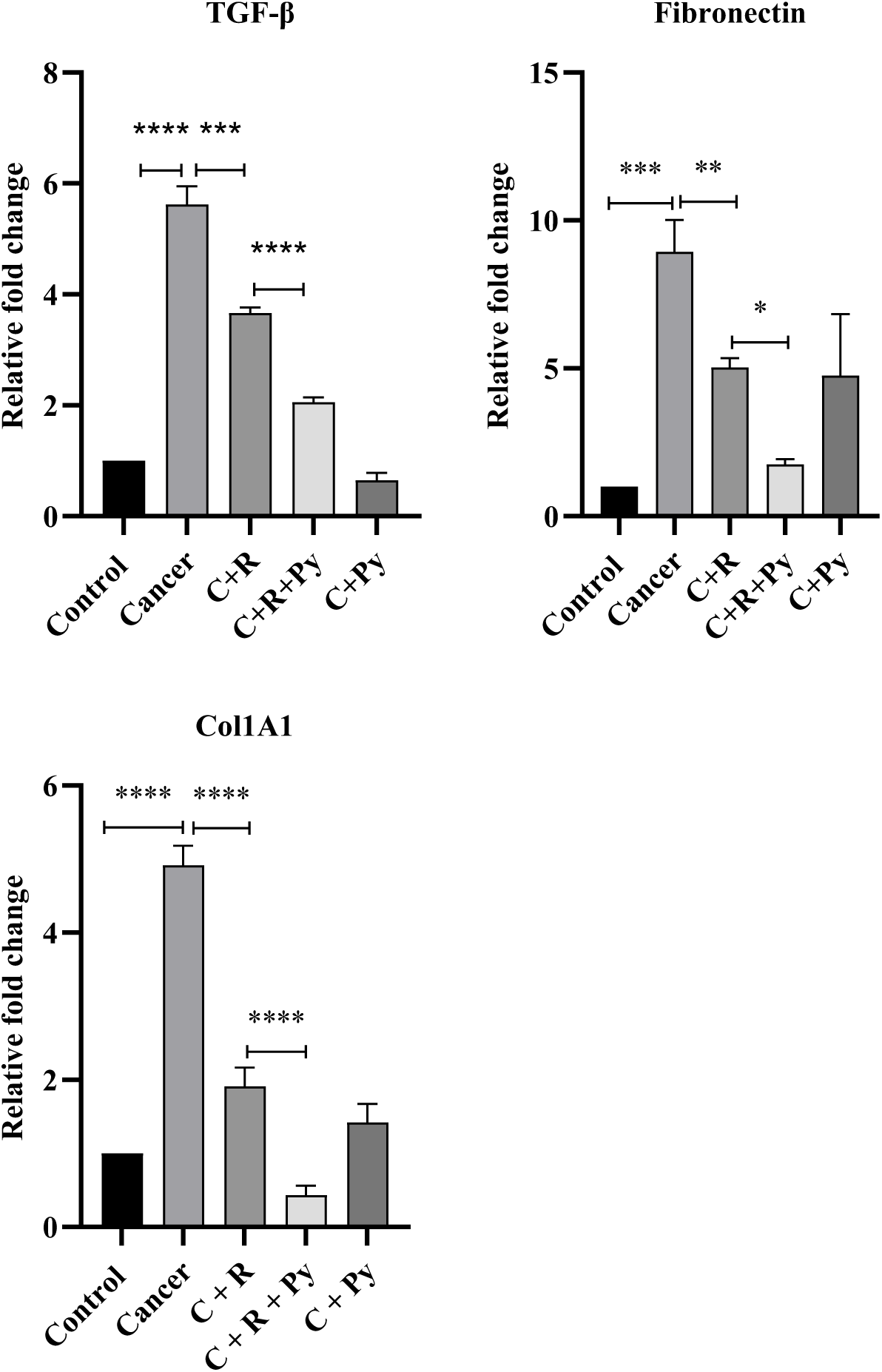
Relative expression profile of host *Tgf-β, Fibronectin* and *Col1A1* respectively across experimental cohorts. (Real-time quantitative PCR analysis showing the relative fold change in *Tgf-β, Fibronectin* and *Col1A1* mRNA expression levels normalized to host reference gene across five experimental groups (n=3 from each group): Control, Cancer (EAC tumour), C+R (Cancer + Radiotherapy), C+R+Py (Cancer + Radiotherapy + Pyrogallol), and C+Py (Cancer + Pyrogallol). Data are presented as mean ± SD; Statistical significance was determined using one-way ANOVA *p < 0.05, **p < 0.01, ***p < 0.001, ****p < 0.0001; ns, not significant).

### 3.5. Pyrogallol enhanced tumor suppressor gene expression in cancer-induced Balb/c mice during radiotherapy

The gene expression levels of the key tumor suppressor genes, *Tp53* and *p21* were analyzed to evaluate the effect of the combination of radiotherapy and pyrogallol in cancer-induced Balb/c mice. In cancer-induced group, the expression of tumor suppressor genes *Tp53* and *p21* was reduced compared with the control group indicating the loss of regulatory mechanism that control cell proliferation and maintain genomic stability during tumor development. The cancer-induced group receiving radiotherapy showed elevated levels of *Tp53* and *p21* genes. The cancer-induced group receiving radiotherapy and pyrogallol showed further significant increase in *Tp53* and *p21* as shown in Figure: 4a and 4b. Mice received only pyrogallol also showed a moderate increase in tumor suppressor gene expression when compared with cancer-induced group.

**Fig 4a, 4b:**
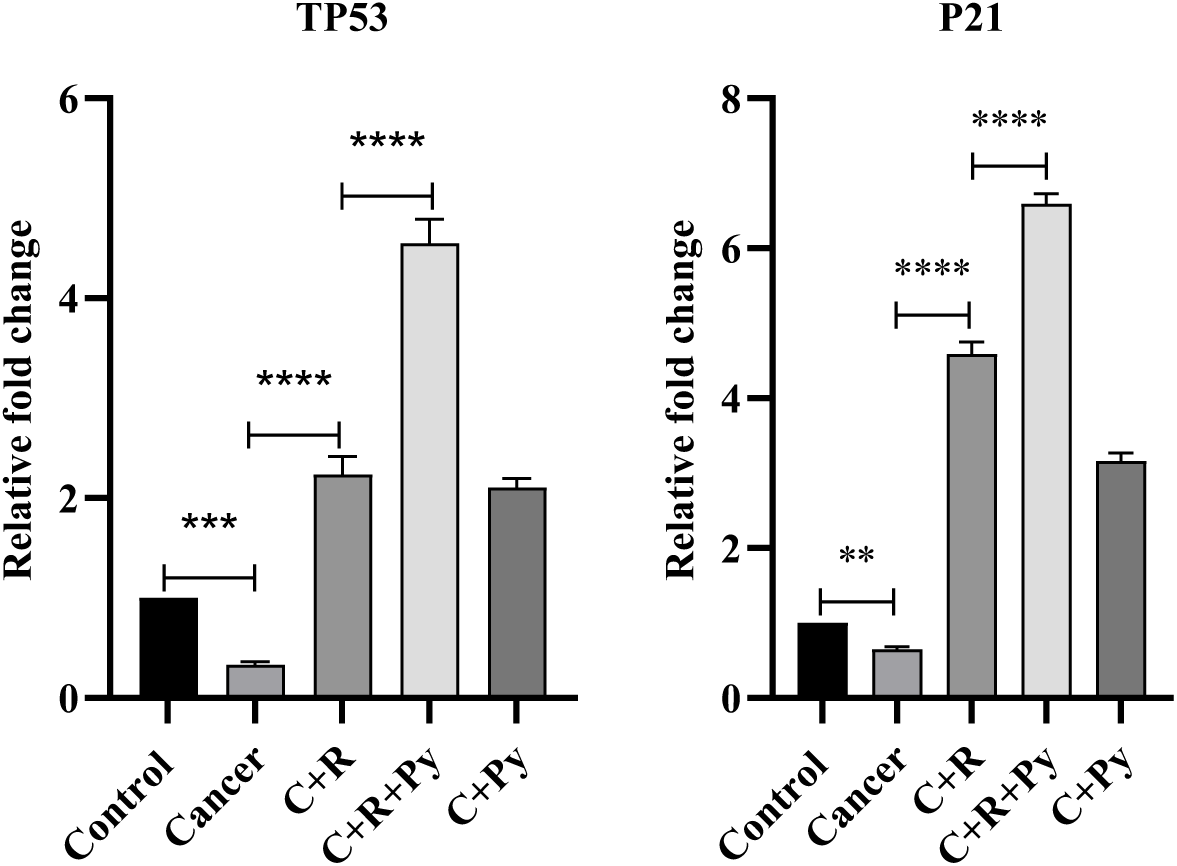
Relative expression profile of host *Tp53* and *p21* respectively across experimental cohorts. (Real-time quantitative PCR analysis showing the relative fold change in *Tp53* and *p21* mRNA expression levels normalized to host reference gene across five experimental groups (n=3 from each group): Control, Cancer (EAC tumour), C+R (Cancer + Radiotherapy), C+R+Py (Cancer + Radiotherapy + Pyrogallol), and C+Py (Cancer + Pyrogallol). Data are presented as mean ± SD; Statistical significance was determined using one-way ANOVA *p < 0.05, **p < 0.01, ***p < 0.001, ****p < 0.0001; ns, not significant).

### 3.6. Pyrogallol inhibits EMT-related gene expression in cancer-induced Balb/c mice during radiotherapy

*N-cadherin* and *Vimentin* are known as canonical markers of the Epithelial-Mesenchymal Transition (EMT), a process that allows stationary cells to become mobile, which is heavily studied in cancer metastasis. *N-cadherin* (neural cadherin) is a transmembrane glycoprotein that facilitates calcium-dependent cell-to-cell adhesion. *Vimentin* is a type III intermediate filament protein that forms part of the cell’s cytoskeleton. The relative gene expression of EMT-related genes *N-cadherin* and *Vimentin* showed increased expression in cancer-induced group compared with the control group. Treatment with radiotherapy significantly reduced the expression of *N-cadherin* and *Vimentin* genes when compared to untreated cancer-induced group suggesting that radiotherapy reduced EMT-related gene expression. Cancer-induced group treated with a combination of radiotherapy and pyrogallol demonstrated further reduction in *N-cadherin* and *Vimentin* gene expression indicating that pyrogallol improved the inhibitory effect of radiotherapy on EMT progression in Figure: 5a and 5b. In mice treated with pyrogallol alone, the expression levels of EMT-related genes were also reduced when compared with the cancer-induced group, although the reduction was less pronounced than that observed in the combined treatment group.

**Fig 5a, 5b:**
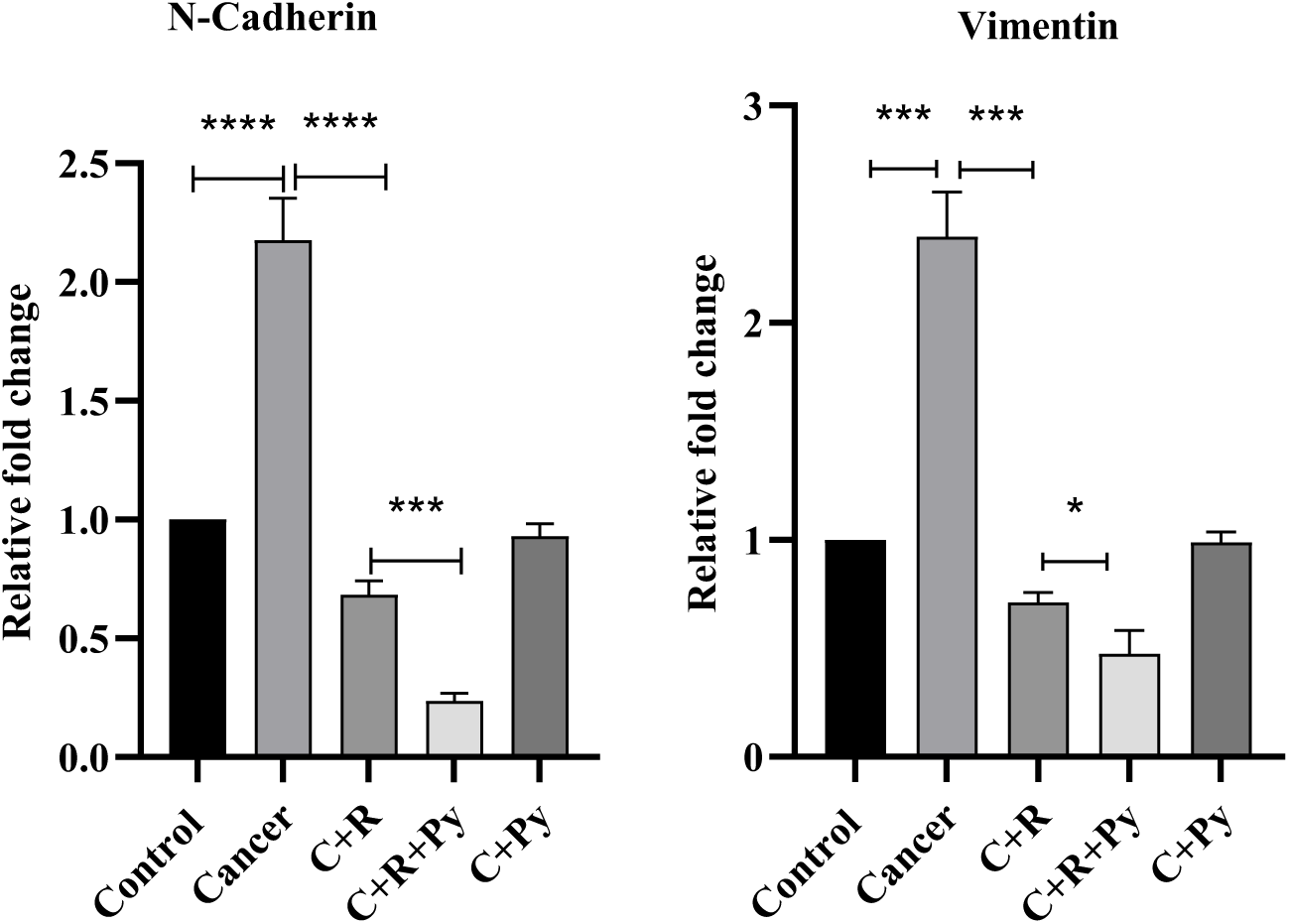
Relative expression profile of host *N-Cadherin* and *Vimentin* respectively across experimental cohorts. (Real-time quantitative PCR analysis showing the relative fold change in *N-Cadherin* and *Vimentin* mRNA expression levels normalized to host reference gene across five experimental groups (n=3 from each group): Control, Cancer (EAC tumour), C+R (Cancer + Radiotherapy), C+R+Py (Cancer + Radiotherapy + Pyrogallol), and C+Py (Cancer + Pyrogallol). Data are presented as mean ± SD; Statistical significance was determined using one-way ANOVA *p < 0.05, **p < 0.01, ***p < 0.001, ****p < 0.0001; ns, not significant).

### 3.7. Pyrogallol attenuates radiation-induced inflammation in cancer-induced Balb/c mice

The expression level of pro-inflammatory cytokines *Il-1α, Il-6* and *Il-12* were evaluated to study radiation-induced inflammation and radiosensitive effect of pyrogallol. The results showed significant up regulation of *Il-1α, Il-6* and *Il-12* in cancer group when compared to control. Treatment with radiation increased these cytokine expressions in radiation group, suggesting increased inflammatory response. However, treatment with pyrogallol along radiation attenuated the cytokine expression significantly indicating the mitigation of radiation-induced inflammatory signaling as depicted in Figure: 6. Additionally, pyrogallol alone group showed reduction in *Il-6* cytokine when compared with control. These findings demonstrated that during cancer pro-inflammatory cytokines such as *Il-1α*, *Il-6* and *Il-12* increased, whereas combined treatment strategy modulated these inflammatory responses.

**Fig 6:**
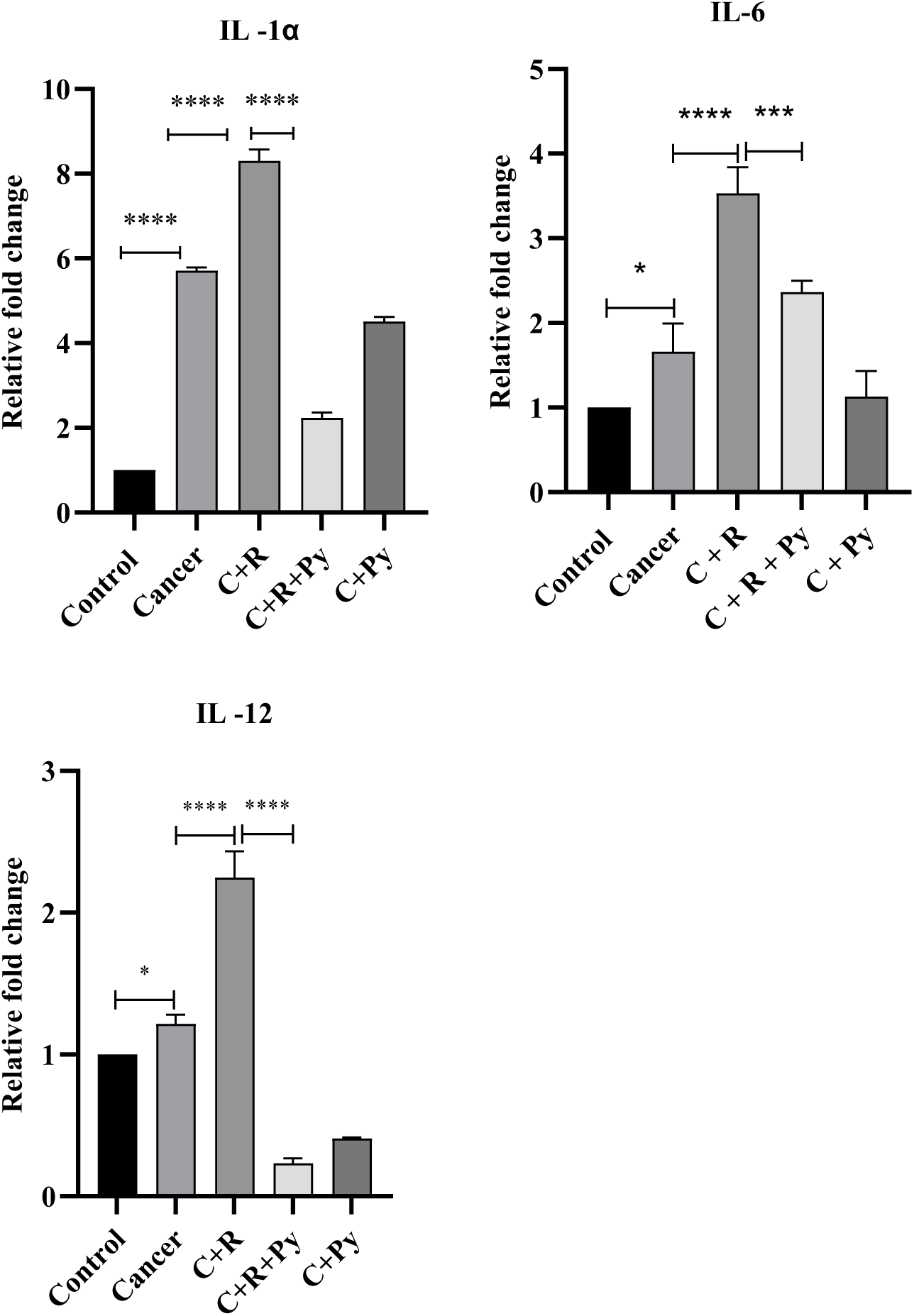
Relative expression profile of host pro-inflammatory cytokines across experimental cohorts. (Real-time quantitative PCR analysis showing the relative fold change in *Il-1α, Il-6, Il-12* mRNA expression levels normalized to host reference gene across five experimental groups (n=3 from each group): Control, Cancer (EAC tumour), C+R (Cancer + Radiotherapy), C+R+Py (Cancer + Radiotherapy + Pyrogallol), and C+Py (Cancer + Pyrogallol). Data are presented as mean ± SD; Statistical significance was determined using one-way ANOVA *p < 0.05, **p < 0.01, ***p < 0.001, ****p < 0.0001; ns, not significant).

### 3.8. Species level abundance identification

To establish a preliminary baseline map of species-level alterations across experimental cohorts, total metagenomic DNA from pooled faecal samples was evaluated using Oxford Nanopore Technology (ONT) whole-metagenome sequencing. As an exploratory pilot survey, these findings reflect representative population-level trends across groups rather than statistically powered individual variances.

Overall, 6 major bacterial phyla were identified across all experimental cohorts. Preliminary observations indicated a gradual contraction of microbial diversity in the treated groups compared to healthy controls at the phylum level, with *Bacillota* and *Bacteroidota* maintaining overall dominance (Fig 7). Relative abundance of top 10 genus across the groups were showed in Fig 8. Unlike standard 16S rRNA gene amplicon profiling, our utilization of whole-genome metagenomic sequencing of pooled faecal samples provided a better resolution of taxonomic mapping down to the species level. Within the *Bacteroides* genus, most identified species exhibited lower relative abundances in the disease and treatment cohorts relative to healthy controls, with two notable exceptions (Fig 9). *Bacteroides caecimuris* and *Bacteroides faecium* increased in cancer induced mice compared to control group (Fig10); studies shown that overgrowth of Bacteroides species may contribute to an immunosuppressive environment that supports tumour growth (17). Furthermore, the opportunistic pathobiont *Acinetobacter baumannii* displayed a notable expansion in the cancer-induced cohort (∼40% increase) and the cancer + radiation group (∼46% increase) compared to healthy controls, whereas its abundance appeared reduced in the drug-treated group. *Blautia coccoides* is a commensal bacterium in the *Lachnospiraceae* family, promotes colonic mucus growth and helps maintain a healthy intestinal barrier by stimulating short-chain fatty acid (SCFA) production also it helps mitigate systemic inflammation (18); decreased gradually across treated cohorts. *Lactiplantibacillus carotarum* is a species of lactic acid bacteria comes under *Bacillota* phylum isolated from fermented carrot juice widely known as probiotic dominates in healthy mice gut while in other 4 treated groups it decreased drastically. *Roseburia intestinalis*, an important butyrate producer that reinforces intestinal integrity and attenuates inflammation (19); alongside commensal *Escherichia coli* strains involved in central metabolic support (20); and *Acutalibacter muris* is generally considered a beneficial or commensal bacterium, all exhibited lower relative representations in the treated groups relative to healthy controls. Coinciding with the loss of protective commensals, *Clostridioides difficile* primarily known as pathogenic bacteria (21), displayed elevated relative abundances in the untreated cancer (∼39% increase) and radiation-only (∼41% increase) cohorts compared to controls. Similarly, *Helicobacter typhlonius* specifically classified as opportunistic pathogen in rodents; overgrowth causes severe inflammation in large intestine lining (22); showed a ∼71% increase in the cancer + radiation cohort. These 10 species abundances were shown individually in Fig 10. *Muribaculum intestinale* known as beneficial gut bacterium which modulates host neurotransmitter synthesis, immune responses, and intestinal barrier function through metabolites such as short-chain fatty acids, succinate, and 3-hydroxybutyrate (23) decreased gradually in cancer and radiation cohort. Likewise, *Muribaculum gordoncarteri,* a key carbohydrate fermenter possessing β-glucosidase, α-arabinase, and α-fucosidase activities (24), which are critical for breaking down plant and host glycans decreased in cancer induced mice compared to healthy mice (Fig 11). *Duncaniella* genus is common mouse gut coloniser within the *Muribaculaceae* family; both the identified species *Duncaniella dubosii* and *Duncaniella sp.*C9 gradually decreased in all the treated groups compared to control group. Interestingly *Duncaniella sp.*C9 completely absent in radiation treated group and also in radiation + drug treated group (Fig 12). Three species under *Phocaeicola* genus; *Phocaeicola dorei*, *Phocaeicola sartorii* and *Phocaeicola vulgatus* commonly known as mutualistic gut commensal that can protect against inflammatory diseases; *P. dorei* and *P. vulgatus* contribute to plant fiber fermentation and SCFA generation, which fuel intestinal cells and strengthen the gut barrier; exhibited lower relative abundances in cancer-bearing mice compared to healthy controls (Fig 13). *Clostridium scindens* is a keystone species that converts primary bile acids into secondary bile acids (like deoxycholic acid), which naturally inhibit the growth and virulence of *Clostridioides difficile* (25) displayed a gradual reduction across all intervention groups, inversely mirroring the observed expansion of *C. difficile*. *Clostridium symbiosum* known as butyrate producer; *Clostridium* sp. C45 and *Clostridium* sp. MD294 promote gut homeostasis by inducing regulatory T (Treg) cells, which help attenuate colonic inflammation and allergic diseases; also produce indole-3-propionic acid (IPA) and butyrate, which strengthen the intestinal mucosal barrier. These three Clostridium species gradually decreased in all the treated groups compared to healthy mice (Fig 14). Among three species under Parabacteroides genus, *Parabacteroides distasonis* found as most abundant; is a beneficial gut bacterium known as a promising next-generation probiotic. It helps decrease weight gain, reduce hyperglycaemia, and lower fat accumulation in the liver also modifies bile acid profiles and boosts succinate production, which activates intestinal gluconeogenesis and improves overall insulin sensitivity (26). *P. distasonis* showed an apparent drop in relative abundance in the untreated cancer (∼22% decrease) and radiation-only (∼16% decrease) groups compared to healthy controls (Fig 15).

**Fig 7:**
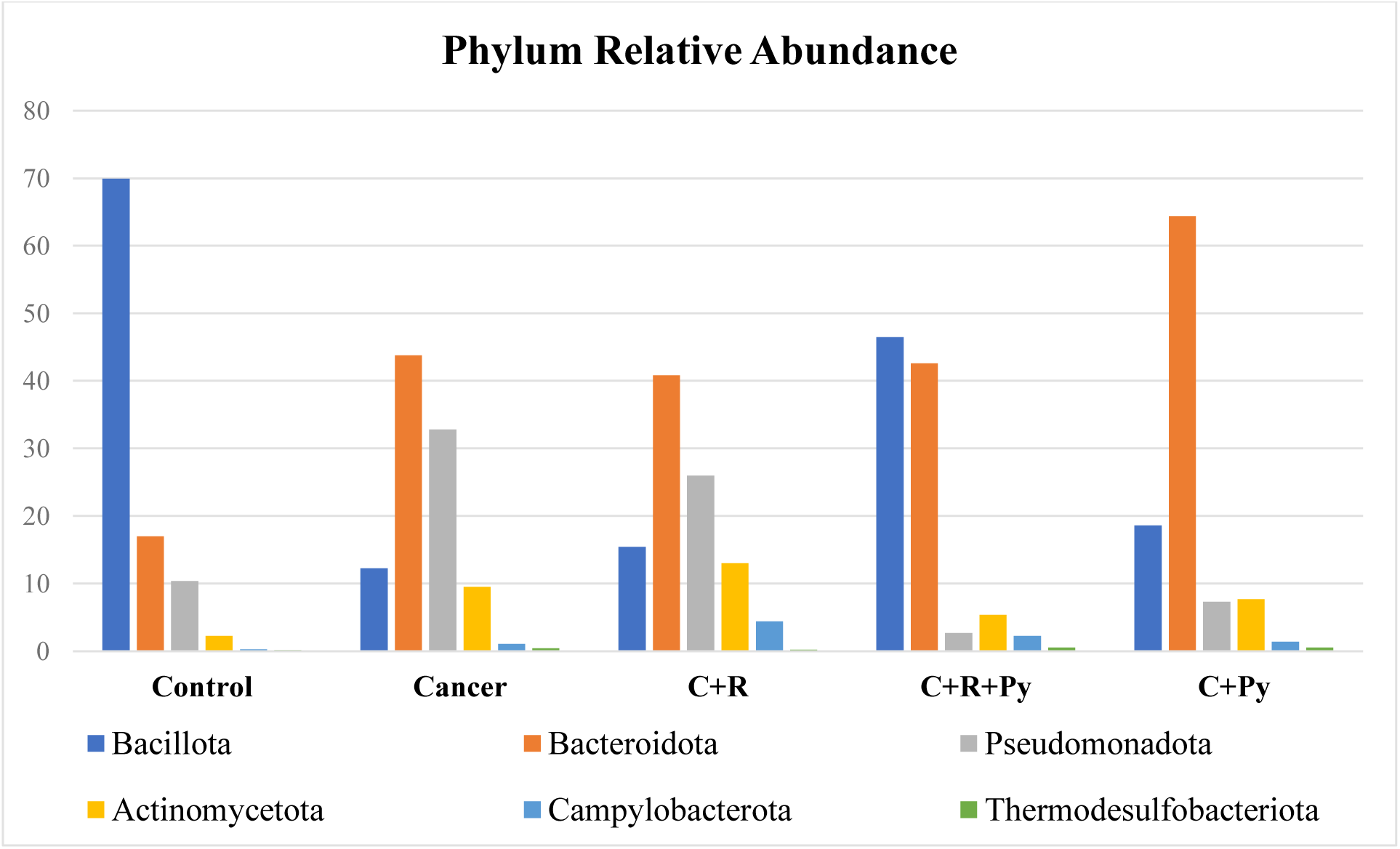
Relative abundance distribution of six Phylum across experimental mouse cohorts. (Bar chart depicting the phylum level sequence abundances of six identified phylum – Bacillota (blue), Bacteroidota (orange), Pseudomonadota (grey), Actinomycetota (yellow), Campylobacterota (sky blue) and Thermodesulfobacteriota (green) which derived from exploratory whole-genome metagenomic sequencing across experimental groups. Values represent baseline species counts obtained from pooled faecal metagenomic profiles (n = 6 mice per cohort pooled samples). [Control: Healthy Control; Cancer: EAC Tumour; C+R: Cancer + Radiotherapy; C+R+Py: Cancer + Radiotherapy + Pyrogallol; C+Py: Cancer + Pyrogallol])

**Fig 8:**
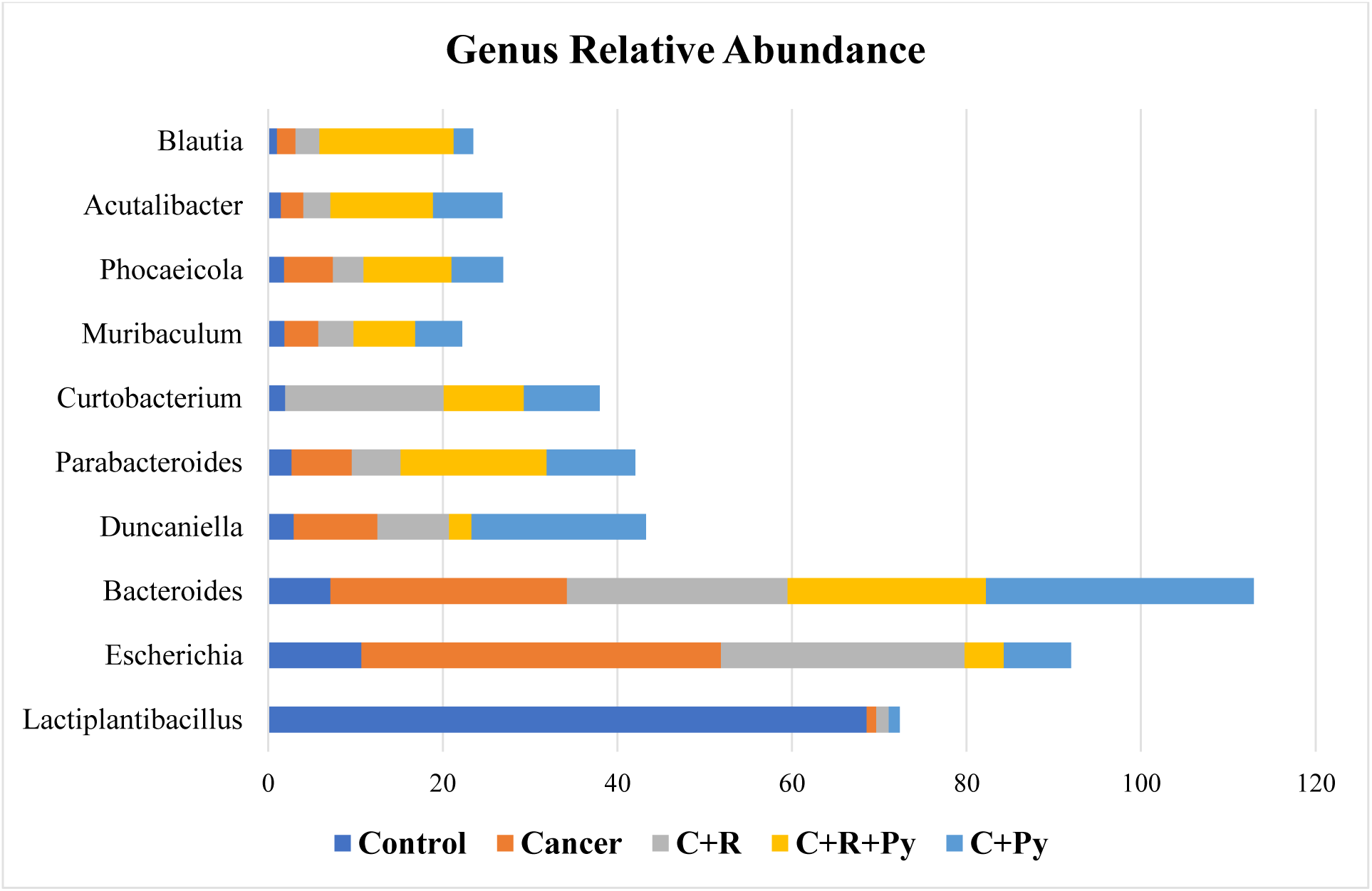
Relative abundance distribution of top 10 genus across experimental mouse cohorts. (Stacked bar chart depicting the top 10 genus level relative sequence abundances from exploratory whole-genome metagenomic sequencing across experimental groups. Values represent baseline species counts obtained from pooled faecal metagenomic profiles (n = 6 mice per cohort pooled samples). [Control: Healthy Control (blue); Cancer: EAC Tumour (orange); C+R: Cancer + Radiotherapy (grey); C+R+Py: Cancer + Radiotherapy + Pyrogallol (yellow); C+Py: Cancer + Pyrogallol (sky blue)])

**Fig 9:**
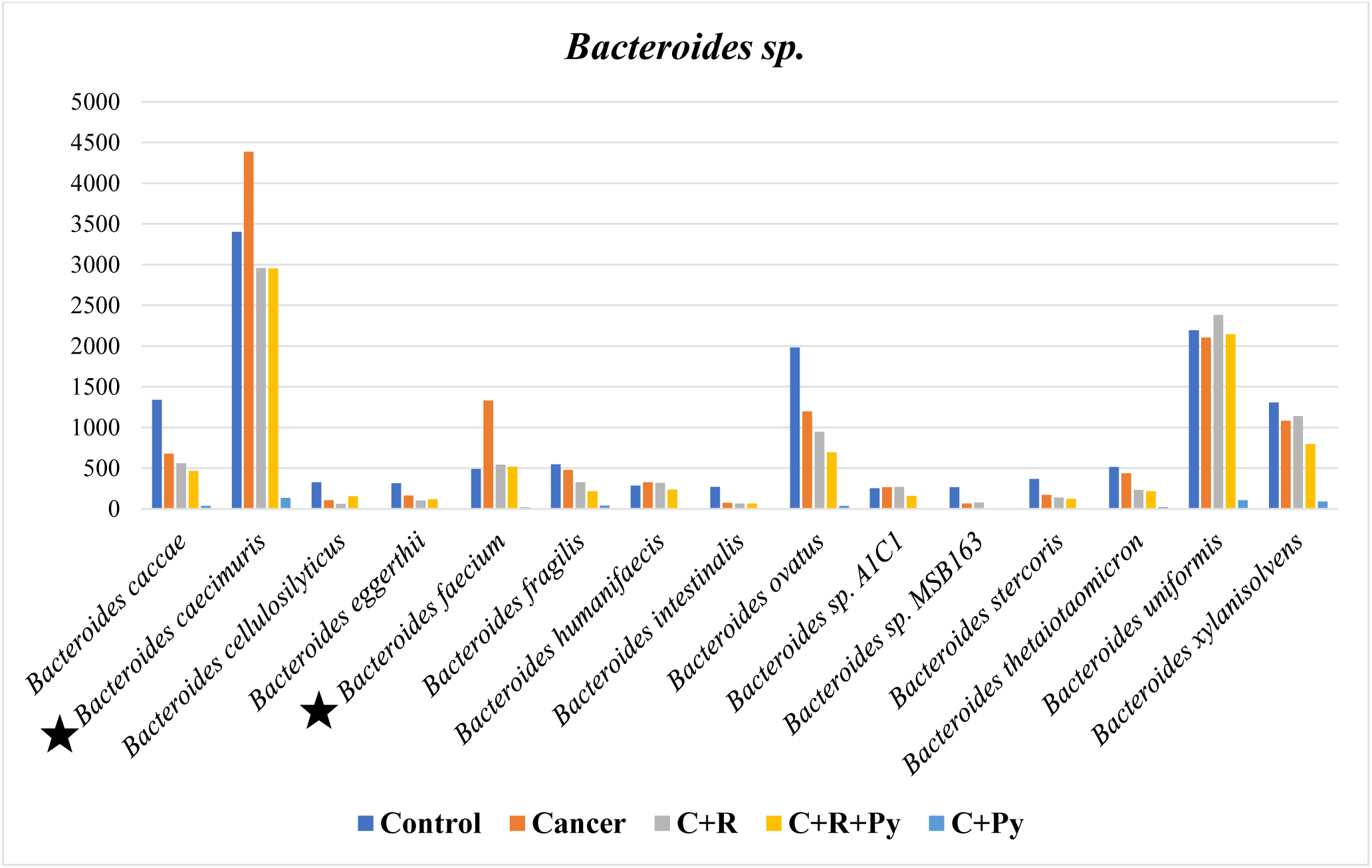
Relative abundance distribution of *Bacteroides species* across experimental mouse cohorts. (Bar chart depicting the species-level sequence abundances of all identified *Bacteroides* species which derived from exploratory whole-genome metagenomic sequencing across experimental groups. Values represent baseline species counts obtained from pooled faecal metagenomic profiles (n = 6 mice per cohort pooled samples). [Control: Healthy Control (blue); Cancer: EAC Tumour (orange); C+R: Cancer + Radiotherapy (grey); C+R+Py: Cancer + Radiotherapy + Pyrogallol (yellow); C+Py: Cancer + Pyrogallol (sky blue)])

**Fig 10:**
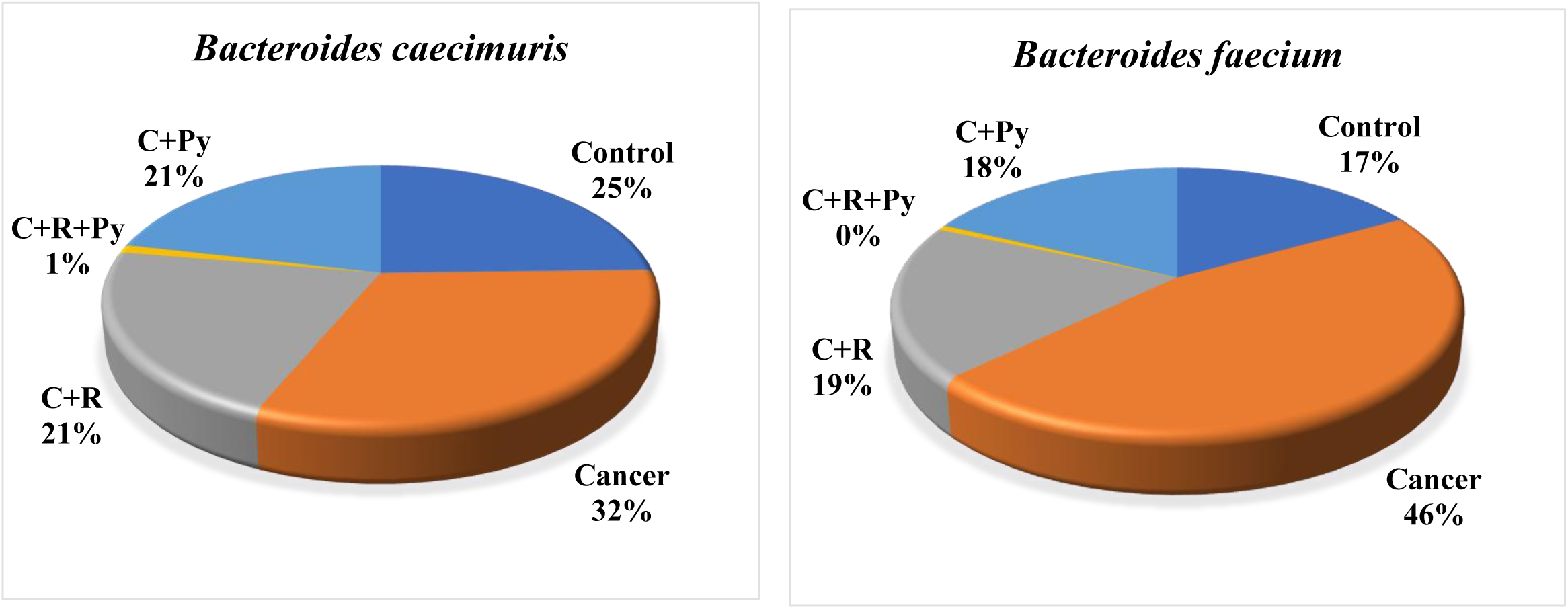
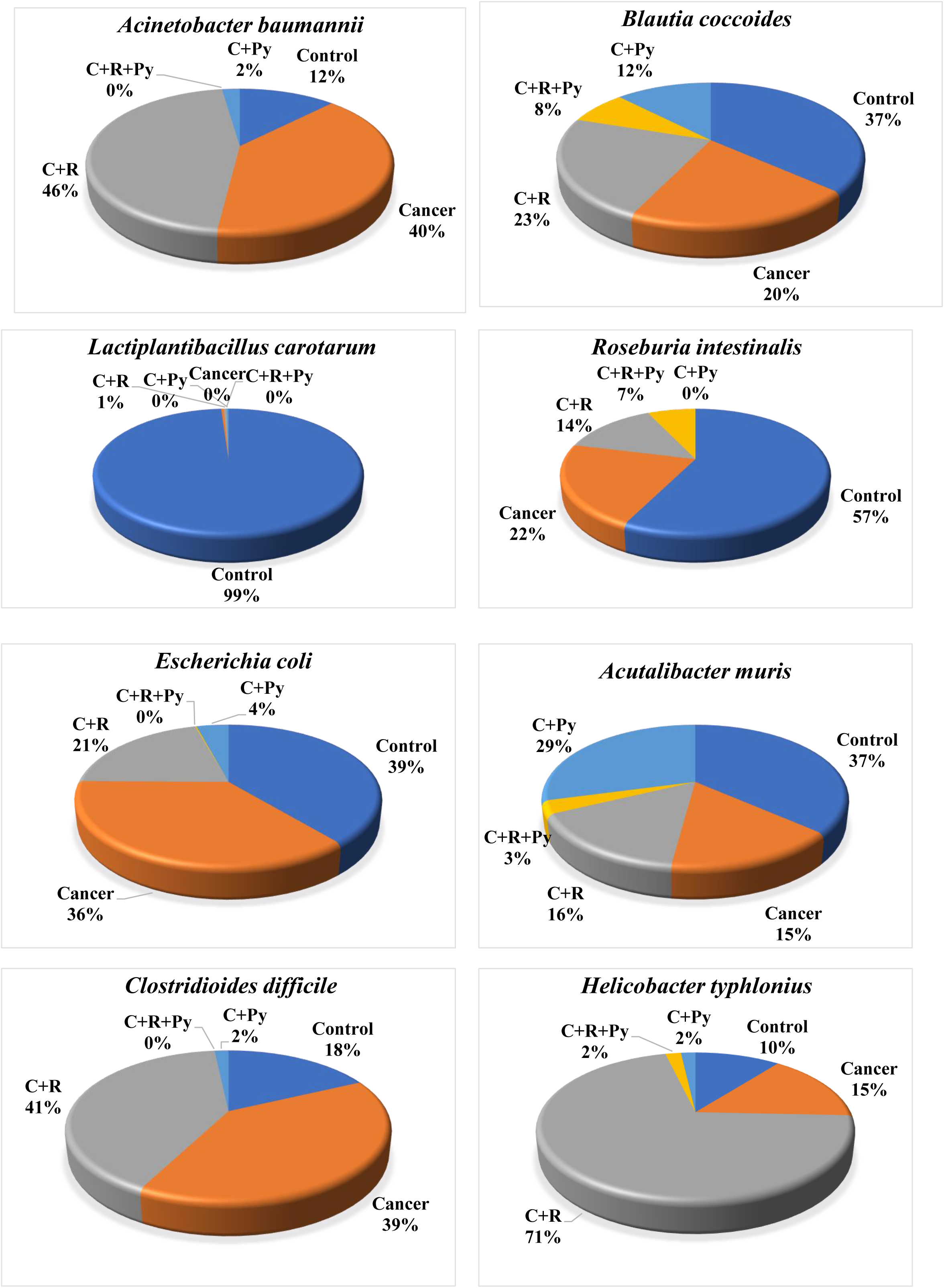
Relative abundance distribution of ten key bacterial species across experimental cohorts. (Pie charts illustrating the relative abundance percentage of ten representative bacterial species— including *Bacteroides caecimuris, Bacteroides faecium, Acinetobacter baumannii, Blautia coccoides, Lactiplantibacillus carotarum*, *Roseburia intestinalis, Escherichia coli, Acutalibacter muris, Clostridioides difficile* and *Helicobacter typhlonius* —across the five experimental cohorts. Abundance percentages were calculated from exploratory whole-genome metagenomic sequencing of pooled faecal samples (*n* = 6 mice per group pooled samples). [Control: Healthy Control; Cancer: EAC Tumor; C+R: Cancer + Radiotherapy; C+R+Py: Cancer + Radiotherapy + Pyrogallol; C+Py: Cancer + Pyrogallol])

**Fig 11:**
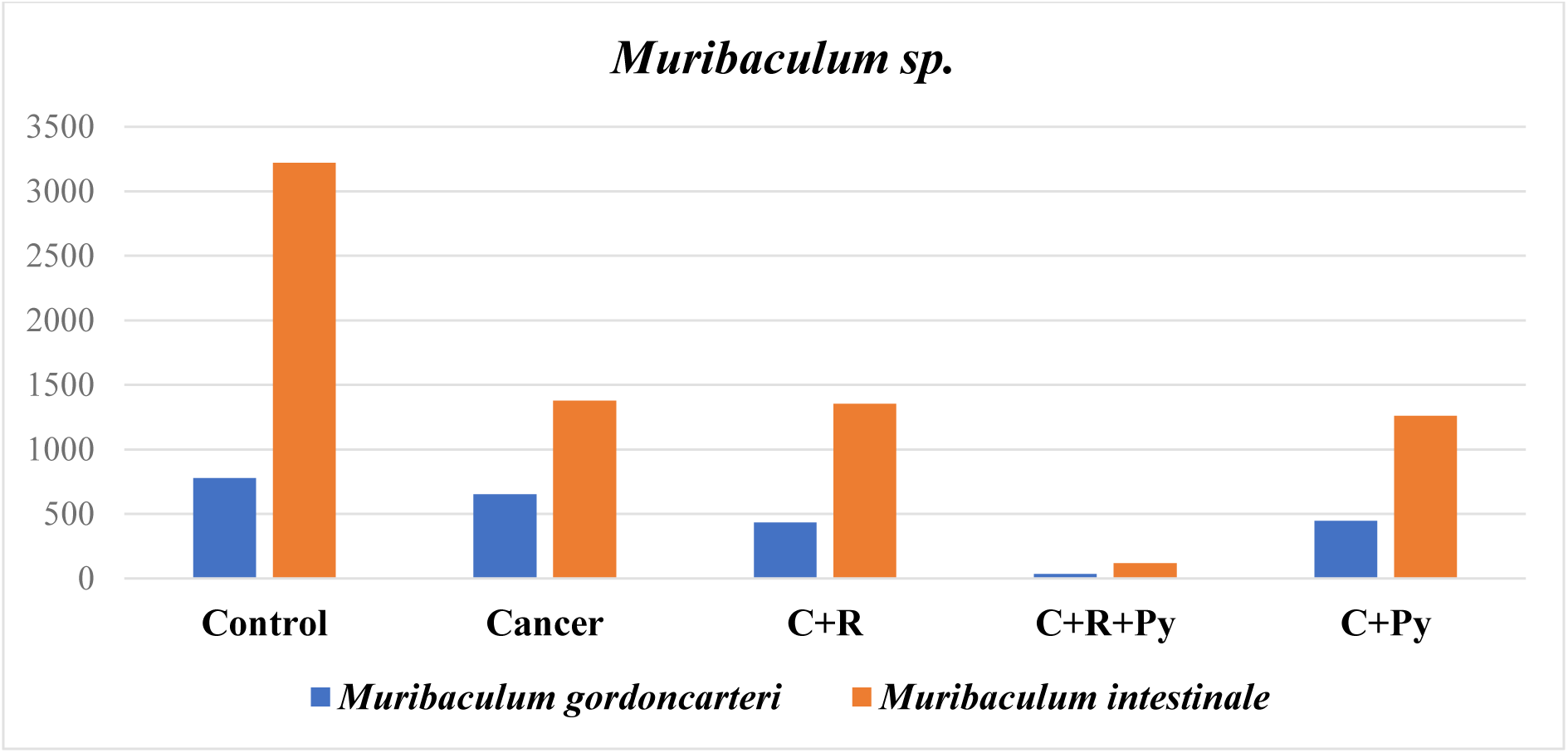
Relative abundance distribution of *Muribaculum species* across experimental mouse cohorts. (Bar chart depicting the species-level sequence abundances of two identified *Muribaculum* species – *Muribaculum gordoncarteri* (blue) and *Muribaculum intestinale* (orange) which derived from exploratory whole-genome metagenomic sequencing across experimental groups. Values represent baseline species counts obtained from pooled faecal metagenomic profiles (n = 6 mice per cohort pooled samples). [Control: Healthy Control; Cancer: EAC Tumour; C+R: Cancer + Radiotherapy; C+R+Py: Cancer + Radiotherapy + Pyrogallol; C+Py: Cancer + Pyrogallol])

**Fig 12:**
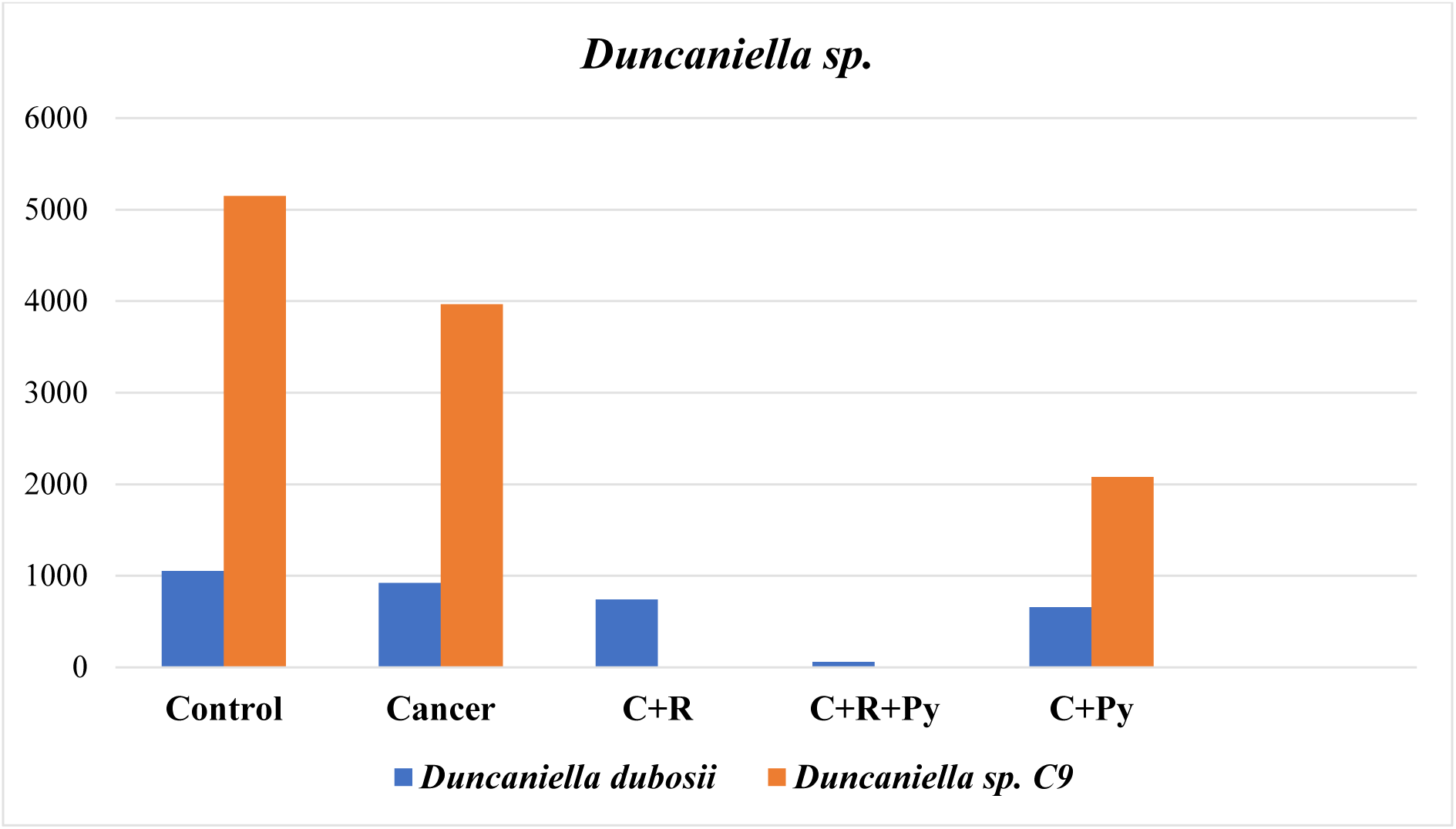
Relative abundance distribution of *Duncaniella* species across experimental mouse cohorts. (Bar chart depicting the species-level sequence abundances of two identified *Duncaniella* species – *Duncaniella dubosii* (blue) and *Duncaniella sp. C9* (orange) which derived from exploratory whole-genome metagenomic sequencing across experimental groups. Values represent baseline species counts obtained from pooled faecal metagenomic profiles (n = 6 mice per cohort pooled samples). [Control: Healthy Control; Cancer: EAC Tumour; C+R: Cancer + Radiotherapy; C+R+Py: Cancer + Radiotherapy + Pyrogallol; C+Py: Cancer + Pyrogallol])

**Fig 13:**
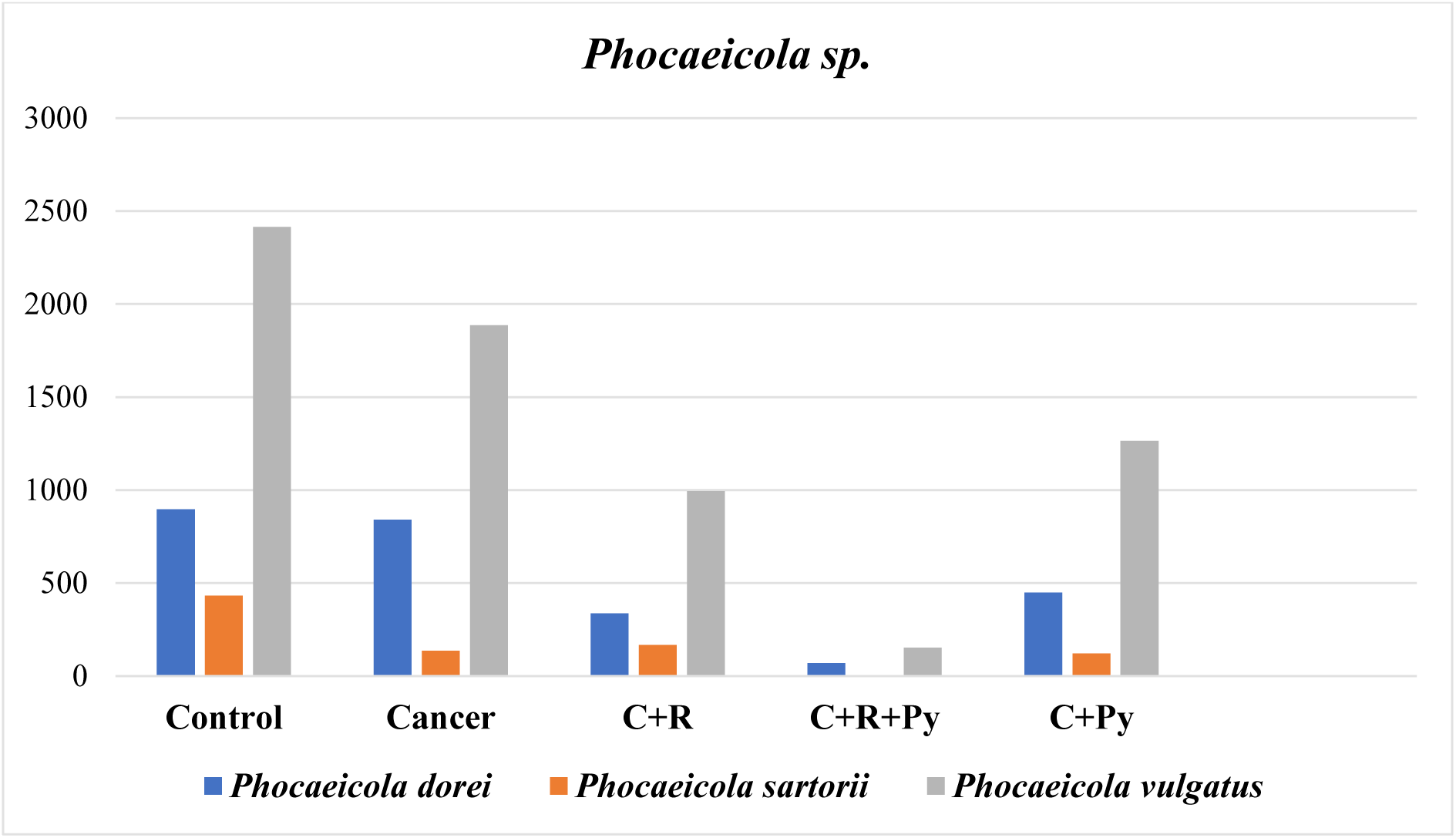
Relative abundance distribution of *Phocaeicola* species across experimental mouse cohorts. (Bar chart depicting the species-level sequence abundances of three identified *Phocaeicola* species – *Phocaeicola dorei* (blue), *Phocaeicola sartorii* (orange) and *Phocaeicola vulgatus* (grey) which derived from exploratory whole-genome metagenomic sequencing across experimental groups. Values represent baseline species counts obtained from pooled faecal metagenomic profiles (n = 6 mice per cohort pooled samples). [Control: Healthy Control; Cancer: EAC Tumour; C+R: Cancer + Radiotherapy; C+R+Py: Cancer + Radiotherapy + Pyrogallol; C+Py: Cancer + Pyrogallol])

**Fig 14:**
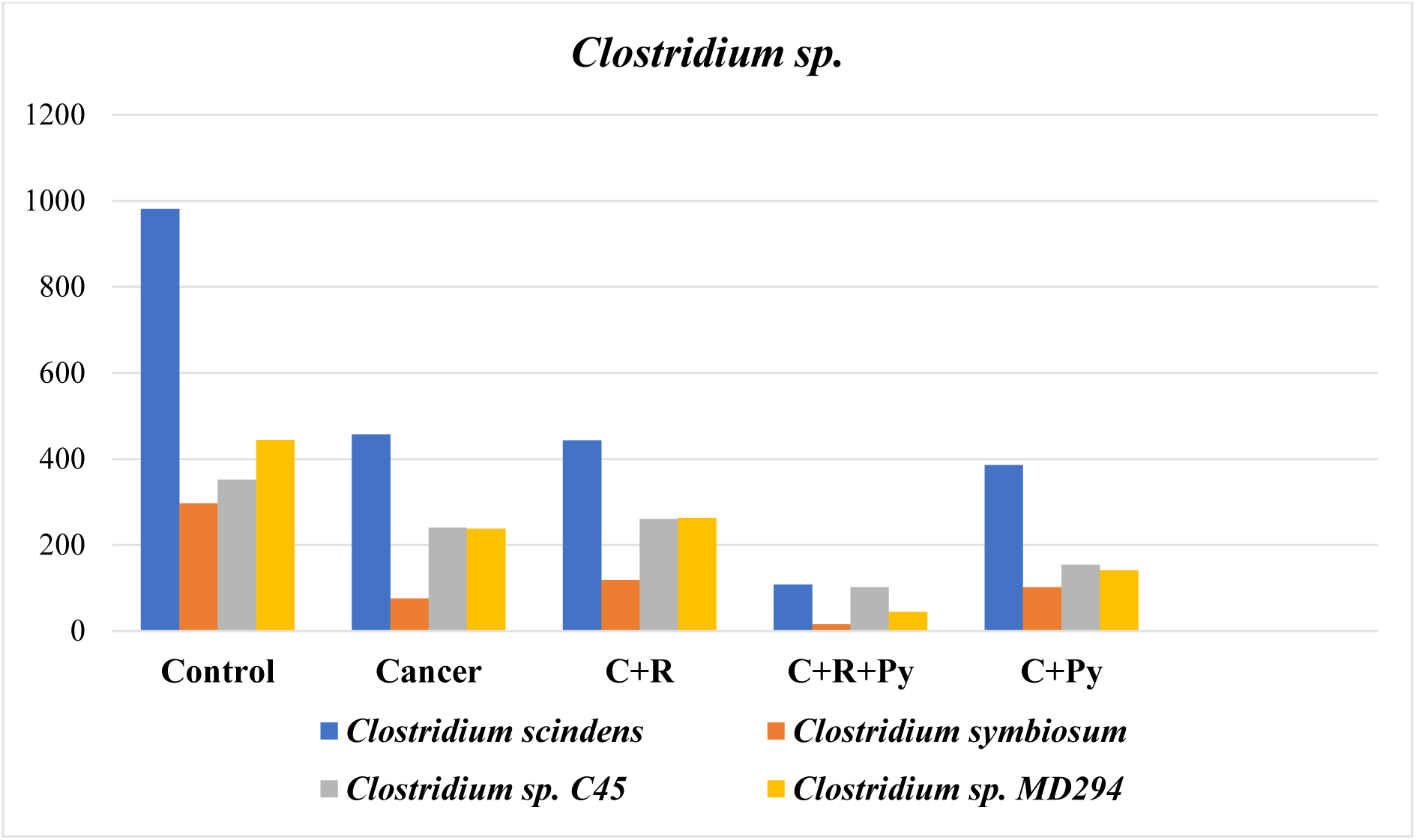
Relative abundance distribution of *Clostridium* species across experimental mouse cohorts. (Bar chart depicting the species-level sequence abundances of four identified *Clostridium* species – *Clostridium scindens* (blue), *Clostridium symbiosum* (orange), *Clostridium sp. C45* (grey) and *Clostridium sp. MD294* (yellow) which derived from exploratory whole-genome metagenomic sequencing across experimental groups. Values represent baseline species counts obtained from pooled faecal metagenomic profiles (n = 6 mice per cohort pooled samples). [Control: Healthy Control; Cancer: EAC Tumour; C+R: Cancer + Radiotherapy; C+R+Py: Cancer + Radiotherapy + Pyrogallol; C+Py: Cancer + Pyrogallol])

**Fig 15:**
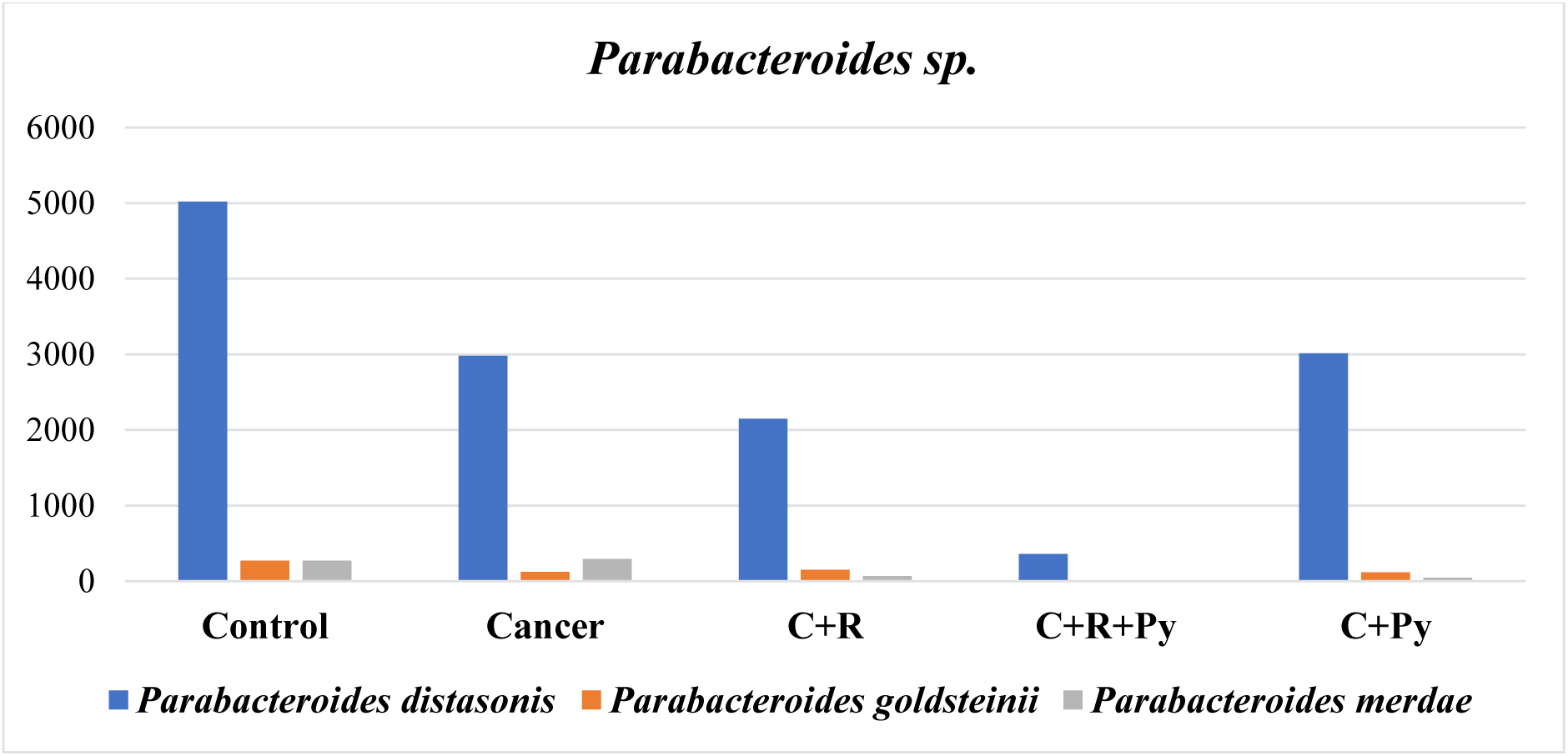
Relative abundance distribution of *Parabacteroides* species across experimental mouse cohorts. (Bar chart depicting the species-level sequence abundances of three identified *Parabacteroides* species - *Parabacteroides distasonis* (blue), *Parabacteroides goldsteinii* (orange), and *Parabacteroides merdae* (grey) which derived from exploratory whole-genome metagenomic sequencing across experimental groups. Values represent baseline species counts obtained from pooled faecal metagenomic profiles (n = 6 mice per cohort pooled samples). [Control: Healthy Control; Cancer: EAC Tumour; C+R: Cancer + Radiotherapy; C+R+Py: Cancer + Radiotherapy + Pyrogallol; C+Py: Cancer + Pyrogallol])

### 3.9. Diversity indices calculation

#### 3.9.1. Alpha diversity

Exploratory alpha diversity metrics suggested a restructuring and numerical contraction of the microbial community following experimental interventions. Observed species richness (Chao1) decreased from 294 unique taxa in healthy controls to 60 in the combination treatment (cancer + radiation + drug treatment) group (C+R+Py). This shift was accompanied by a rise in the Berger-Parker index (measured for species dominance) to 0.94, pointing toward an ecological restructuring favouring dominant surviving taxa under combined exposure, alongside a lower overall community evenness (Shannon index = 0.41) and lower community evenness and richness (Simpson’s index = 0.11) in C+R+Py (Fig 16).

**Fig 16:**
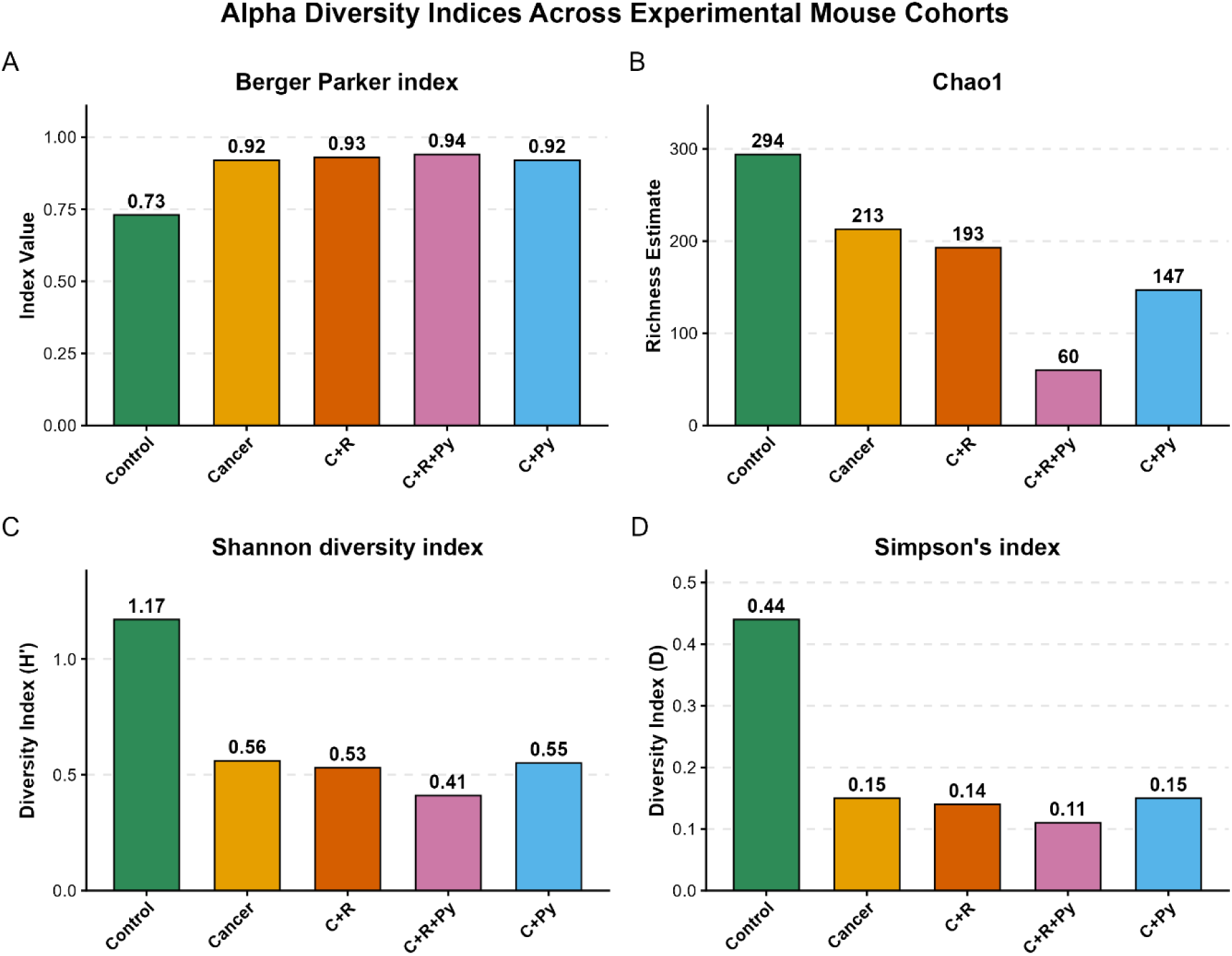
Alpha diversity metrics of murine gut bacterial communities across experimental cohorts. (Comparison of species-level gut microbial alpha diversity derived from exploratory whole-genome metagenomic sequencing across five experimental groups: (A) Berger-Parker dominance index, reflecting the relative numerical dominance of the most abundant taxon; (B) Chao1 richness estimate, indicating observed species richness; (C) Shannon diversity index (H’), accounting for both species richness and evenness; and (D) Simpson’s diversity index (D). Bar values represent representative cohort estimates calculated from pooled faecal samples (n = 6 mice per group pooled samples). [Control: Healthy Control; Cancer: EAC Tumor; C+R: Cancer + Radiotherapy; C+R+Py: Cancer + Radiotherapy + Pyrogallol; C+Py: Cancer + Pyrogallol])

### Beta diversity

Principal Coordinate Analysis (PCoA) based on Bray-Curtis dissimilarity illustrated qualitative shifts in global community structures across cohorts. The primary axis (PCoA 1, accounting for 61.9% of total variance) separated the healthy control cohort from all intervention groups. The untreated cancer and radiation-only (C+R) cohorts clustered in relative proximity, suggesting broadly comparable overall profile configurations under disease and radiation stress. In contrast, the combination treatment cohort (C+R+Py) migrated along a distinct trajectory along PCoA 2 (29.3% variance), pointing toward a unique, specialized ecological restructuring under combined drug and radiation exposure. (Fig 17).

**Fig 17:**
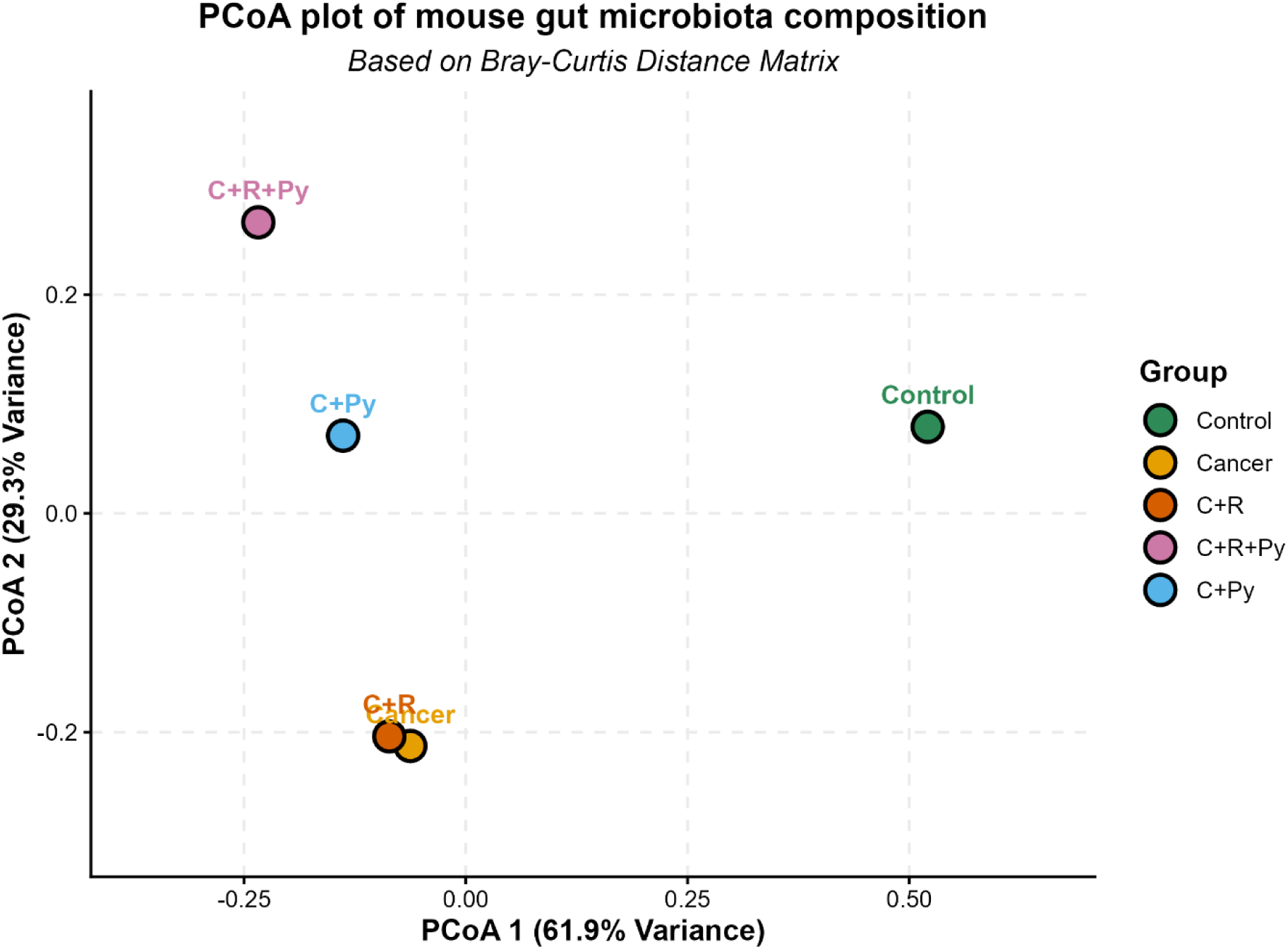
Principal Coordinate Analysis (PCoA) plot of mouse gut microbiota beta diversity across experimental cohorts based on Bray-Curtis dissimilarity. (Ordination plot based on the Bray-Curtis dissimilarity matrix derived from exploratory whole-genome metagenomic sequencing of relative species abundances. The primary axis (PCoA 1) accounts for 61.9% of the total compositional variance, separating the healthy Control group from disease and treatment cohorts. The secondary axis (PCoA 2) accounts for 29.3% of the variance, illustrating distinct community trajectories for the pyrogallol-treated cohorts (C+Py and C+R+Py) relative to the untreated Cancer and radiation-only (C+R) cohorts. Each point represents a representative pooled cohort profile (*n* = 6 mice per group pooled samples). [Control: Healthy Control; Cancer: EAC Tumour; C+R: Cancer + Radiotherapy; C+R+Py: Cancer + Radiotherapy + Pyrogallol; C+Py: Cancer + Pyrogallol])

### Spearman’s Correlation

Integrated Spearman’s rank correlation analysis was performed to explore potential baseline alignments between host intestinal molecular expression profiles and gut bacterial species trends:

### Apoptotic pathways

Anti-apoptotic gene *Bcl2* positively correlated with the relative abundance of *Bacteroides caecimuris* and *Bacteroides faecium* whose abundance increased in cancer mice as we discussed earlier. In contrast, pro-apoptotic genes *Bax*, *Casp3* and *Casp7* exhibited negative correlation trends with these same *Bacteroides* species (Fig 18).

**Fig 18:**
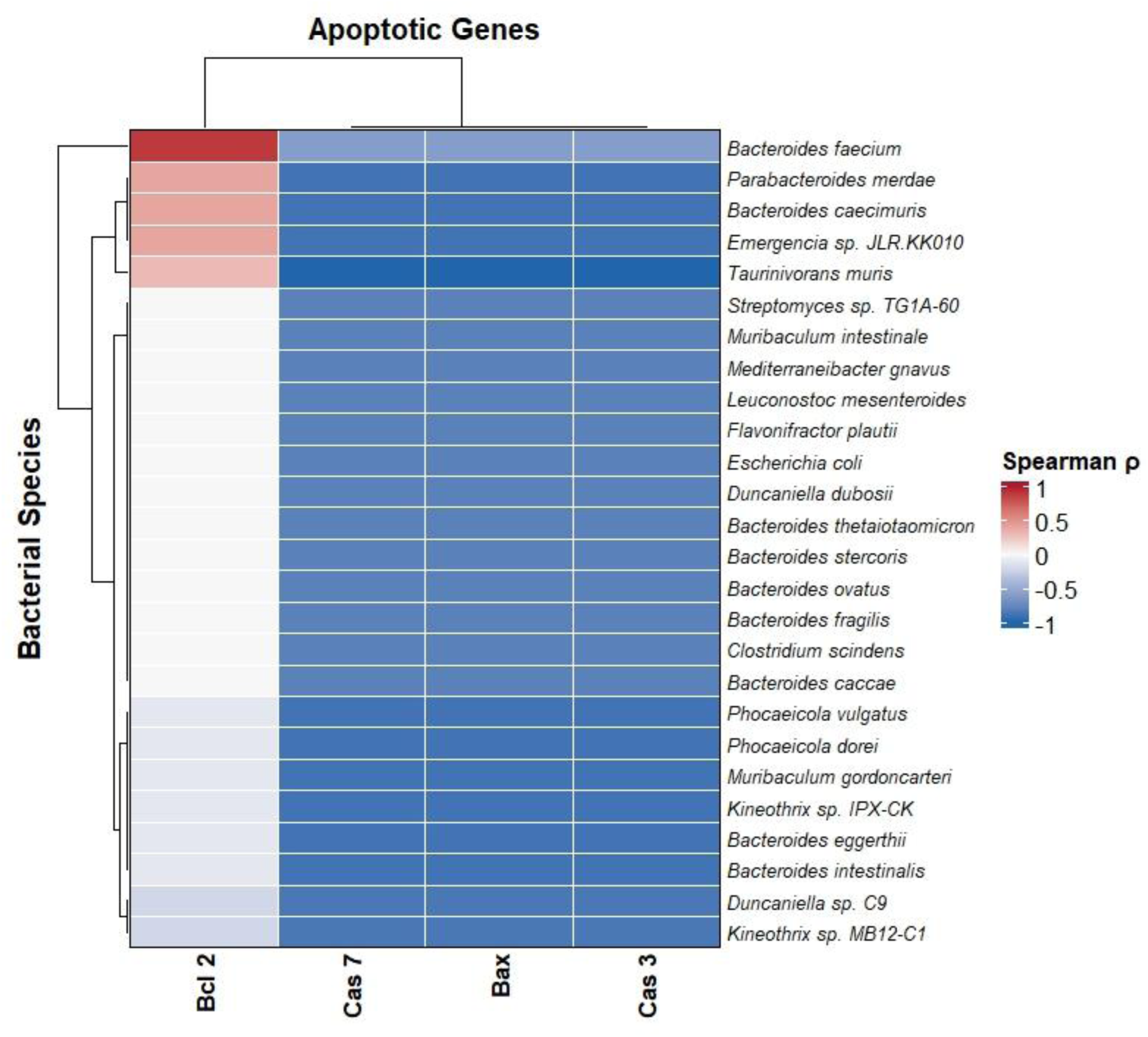
Hierarchical clustering heatmap of Spearman’s rank correlation between host apoptotic gene expression profiles and bacterial species relative abundances. (Correlation heatmap between host apoptotic genes expression and relative abundance of gut microbiome was made using normalized z-score for both the dataset. The colour gradient illustrates Spearman’s rank correlation coefficient (ρ), ranging from positive correlations (red, ρ = +1) to negative correlations (blue, ρ = -1). Both host genes (x-axis): Anti-apoptotic gene *Bcl2* and pro-apoptotic genes *Bax*, *Casp3* and *Casp7* and bacterial species (y-axis) are hierarchically clustered based on Euclidean distance and average linkage to display co-association patterns across experimental cohorts)

### Inflammatory cascades

Inflammatory cytokines *Il-1α* and *Il-12* positively correlated with *Bacteroides faecium, Acinetobacter baumannii,* and two other species whose abundance increased in cancer mice compared to healthy mice. Similarly, *Il-6* negatively associated with commensals such as *Muribaculum gordoncarteri*, *Phocaeicola dorei*, *Phocaeicola vulgatus*, *Bacteroides eggerthii, Bacteroides intestinalis, Parabacteroides distasonis* whose abundance decreased in cancer mice compared to healthy mice (Fig 19).

**Fig 19:**
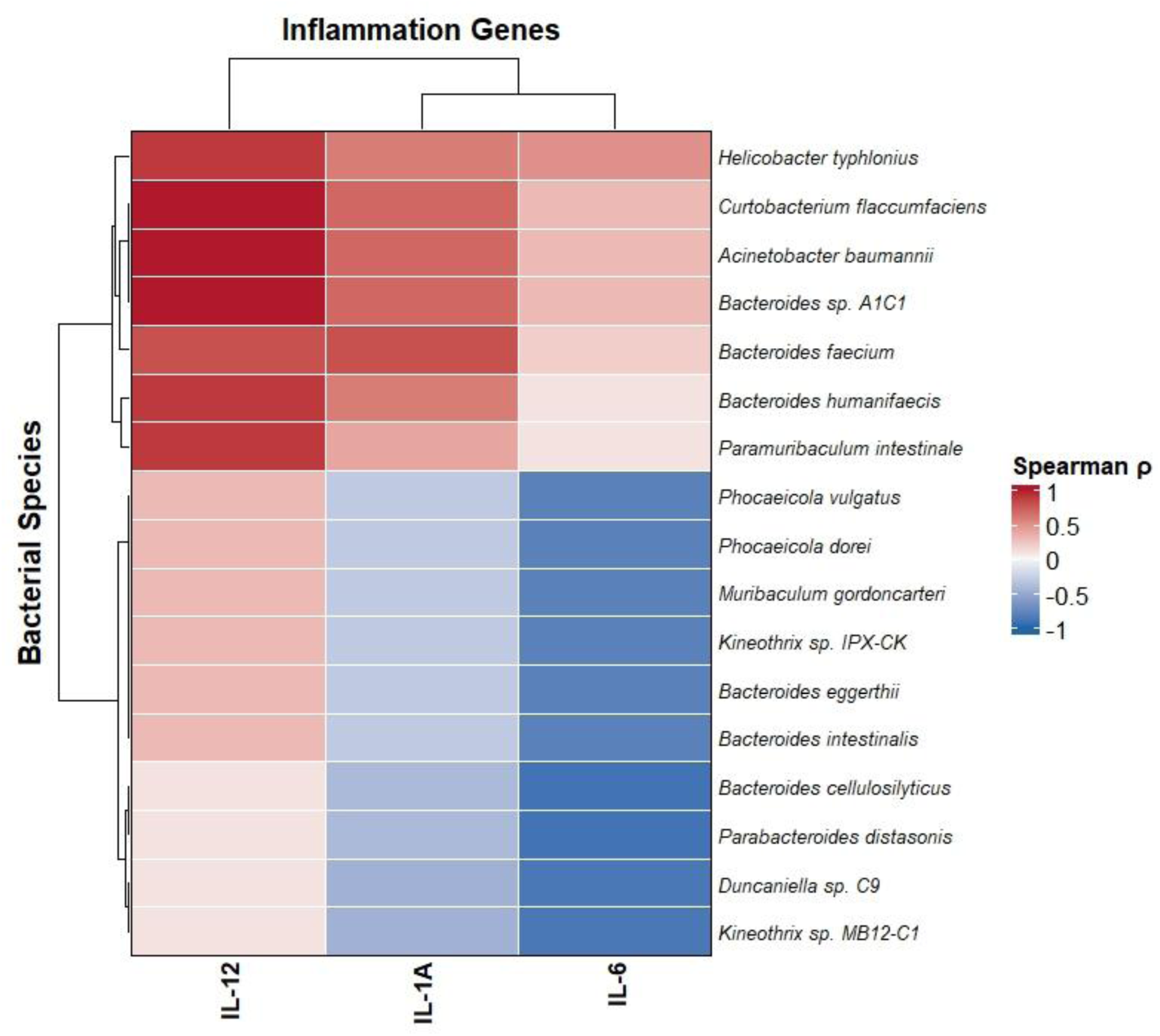
Hierarchical clustering heatmap of Spearman’s rank correlation between host inflammatory gene expression profiles and bacterial species relative abundances. (Correlation heatmap between inflammatory genes expression and relative abundance of gut microbiome was made using normalized z-score for both the dataset. The colour gradient illustrates Spearman’s rank correlation coefficient (ρ), ranging from positive correlations (red, ρ = +1) to negative correlations (blue, ρ = -1). Both host genes (x-axis): inflammatory cytokines (*Il-12, Il-1α, Il-6*) and bacterial species (y-axis) are hierarchically clustered based on Euclidean distance and average linkage to display co-association patterns across experimental cohorts)

### Oncogenic, EMT and Fibrotic drivers

Tumour suppressor gene *Tp53* and *p21* negatively correlated with pathobiont associated clusters including *Bacteroides caecimuris*, *Bacteroides faecium, Acinetobacter baumannii* whose abundance increased in cancer mice compared to healthy mice. Conversely, key regulators of fibrosis and ECM remodelling (*Tgf-β*, *Col1A1*, and *Fibronectin*), cell cycle progression (*Cdk4*), proliferative markers (*Pcna*), and EMT markers (*N-cadherin* and *Vimentin*) shared positive correlation trends with these species, aligning with their co-elevation in cancer-induced cohorts. (Fig 20).

**Fig 20:**
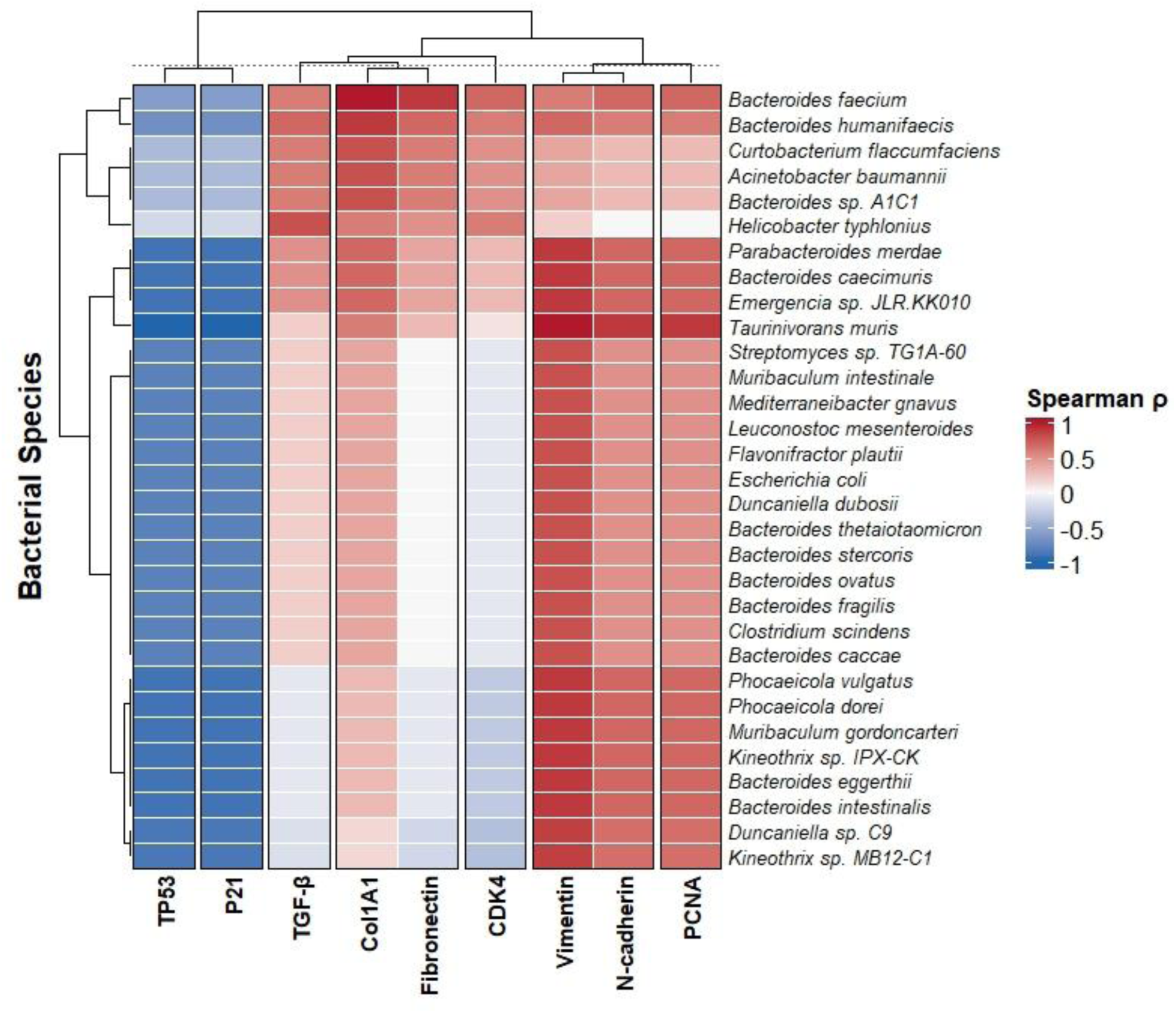
Hierarchical clustering heatmap of Spearman’s rank correlation between host intestinal gene expression profiles and bacterial species relative abundances. (Correlation heatmap between various genes expression and relative abundance of gut microbiome was made using normalized z-score for both the dataset. The colour gradient illustrates Spearman’s rank correlation coefficient (ρ), ranging from positive correlations (red, ρ = +1) to negative correlations (blue, ρ = -1). Both host genes (x-axis): tumour suppressors [*Tp53*, *p21*], fibrotic markers [*Tgf-β*, *Col1A1*, *Fibronectin*], cell cycle regulator [*Cdk4*], EMT markers [*Vimentin*, *N-cadherin*], and proliferation marker [*Pcna*] and bacterial species (y-axis) are hierarchically clustered based on Euclidean distance and average linkage to display co-association patterns across experimental cohorts)

## 4. Discussion

### Pyrogallol Enhances Radiation-Induced Apoptosis and Tumour Suppression Amidst Microbial Diversity Contraction

Ionizing radiation inflicts cellular damage by generating reactive oxygen species (ROS) and inducing DNA strand breaks, ultimately triggering apoptotic cascades in malignant tissues. In the present study, exposing Ehrlich ascites carcinoma (EAC) to localized ionizing radiation (8 Gy LINAC) significantly upregulated host pro-apoptotic markers (*Bax, Casp3, Casp7*) and downregulated the anti-apoptotic oncogene *Bcl2*, confirming the activation of classic radiation-induced programmed cell death. This apoptotic response was further underpinned by the elevated expression of tumour suppressors *Tp53* and *p21*, indicating a functional, p53-mediated apoptotic axis.

Crucially, co-treatment with pyrogallol and radiotherapy synergistically amplified this signalling cascade, driving further suppression of *Bcl2* and robust upregulation of pro-apoptotic and tumour suppressor genes. This confirms that pyrogallol acts as an effective radio-sensitizing agent. These findings align with previous phytochemical-based radiosensitization literature; for instance, Abdraboh et al. (27) demonstrated that a breast safeguard phytochemical cocktail significantly enhanced *p53* and *Bax* while suppressing *Bcl2* to drive apoptosis in carcinoma cells, and Wang et al. (28) reported a similar apoptotic synergy using resveratrol in glioblastoma models.

Parallel to these host-level molecular shifts, high-resolution whole-genome metagenomic sequencing on the ONT MinION platform provided an exploratory map of structural shifts within the murine gut bacteriome. The aggressive combination of tumour progression, tissue-damaging ionizing radiation, and xenobiotic clearing of pyrogallol coincided with an apparent contraction in alpha diversity, where species richness collapsed from 294 unique taxa in healthy controls to just 60 species in the combined treatment group (C+R+Py). This collapse was characterized by a drop in the Shannon diversity index (0.41) and an escalation of the Berger-Parker dominance index to 0.94. Similar patterns of microbial depletion have been observed in clinical settings; for instance, Pyrosequencing analysis of 16S rRNA gene revealed that overall gut microbial alpha diversity has been depleted in the patients undergoing pelvic cancer radiotherapy (29).

While such diversity shifts often reflect ecological distress, taxonomic assignment suggested that this altered niche was 94% dominated by *Parabacteroides distasonis* in C+R+Py group, a well-documented, next-generation probiotic strain. Principal Coordinate Analysis (PCoA) supported this unique restructuring; while the untreated cancer and radiation-only (C+R) cohorts clustered closely together in a shared dysbiotic state, the C+R+Py group migrated along a distinct trajectory (PCoA 2: 29.3% variance). These exploratory trends suggests that pyrogallol co-treatment may foster a specialized ecological environment that favours the relative expansion of this protective obligate anaerobe.

Spearman’s rank correlation mathematically validated this host-microbe alignment: host tumour suppressors (*Tp53, p21*) coupled negatively with pathogenic clusters, while the anti-apoptotic *Bcl2* gene correlated positively with opportunistic tumour-supporting strains like *Bacteroides caecimuris* and *Bacteroides faecium*. These preliminary associations indicate that pyrogallol-mediated tumour suppression parallels a gut landscape largely cleared of opportunistic pathogens.

### Suppression of Anti-Proliferative, EMT, and Fibrotic Drivers Modulates the Gut Pathobiome

Malignant progression relies heavily on escaping cell cycle regulation, activating epithelial-mesenchymal transition (EMT) for invasion, and inducing fibrotic tissue remodelling. Cyclin-dependent kinases, particularly host *Cdk4*, act as the primary regulatory engines driving this cell cycle progression. In tumour-bearing mice, *Cdk4* and Proliferating Cell Nuclear Antigen (*Pcna*) were heavily upregulated, signifying rapid tumour cell proliferation. While radiation alone moderately reduced these markers, pyrogallol co-treatment caused a further decline in both *Cdk4* and *Pcna* expression, effectively putting the brakes on cancer cell proliferation. This matches the findings of Xu et al. (2017), where the polyphenol diosmetin acted as a radiosensitizer in A549 cells, downregulating CDKs to induce a tight G1-phase cell cycle arrest (30).

Simultaneously, pyrogallol enhanced the suppressive effects of radiotherapy on metastatic and tissue-remodelling pathways, downregulating key EMT markers (*N-cadherin, Vimentin*) and critical regulators of fibrosis (*Tgf-β, Col1A1, Fibronectin*). This multi-target suppression is consistent with Kaushik et al. (2017), who demonstrated that ionizing radiation restricts mesenchymal progression by reducing vimentin (31), and Park et al. (2022), who showed that the clinical inhibitor vactosertib successfully halts radiation-induced extracellular matrix deposition by blocking *Tgf-β* signalling (32).

On the microbial side, this down-regulation of host proliferative and structural remodelling genes directly mirrored a major species-level tug-of-war between opportunistic pathogens and health promoting commensals in the gut ecosystem. Under the untreated cancer and radiation-only states, opportunistic pathobiont *Acinetobacter baumannii* surged by up to 46%. *A. baumannii* is known to exploit compromised mucosal barriers and aggravate systemic inflammatory stress. Interestingly, pyrogallol administration completely reversed this pathogenic bloom, outcompeting these opportunistic strains by fueling the massive 94% expansion of *P. distasonis* in C+R+Py.

Spearman’s rank correlation suggested a tight mechanical link between these host and microbial alterations: *Cdk4, Pcna*, EMT drivers (*N-cadherin, Vimentin*), and the tissue-fibrosis axis (*Tgf-β, Col1A1, Fibronectin*) all shared robust positive correlations with *B. caecimuris, B. faecium* and *A. baumannii*. Because the overgrowth of *B. caecimuris* and *B. faecium* is known to foster an immunosuppressive, pro-proliferative microenvironment, pyrogallol’s ability to simultaneously suppress host oncogenic signalling and eliminate these pathobionts highlights its dual-action therapeutic potential.

### Attenuation of Radiation-Induced Inflammation and Restoration of Colonization Resistance

One of the most clinically challenging side effects of radiotherapy is localized and systemic inflammation, driven by the surge of pro-inflammatory cytokines. In this study, localized 8 Gy LINAC irradiation sharply upregulated intestinal expression of *Il-1α, Il-6*, and *Il-12*, indicating severe radiation-induced tissue inflammation. However, pyrogallol co-treatment dramatically attenuated these cytokine expressions, demonstrating a prominent anti-inflammatory profile. This protective effect mirrors in a study, where the administration of 4-methylumbelliferone significantly reduced radiation-induced *Il-1α* and *Il-6* expression in fibrosarcoma models, a mechanism suggested to minimize cellular stress and improve overall therapeutic survival (33).

This molecular anti-inflammatory profile closely aligns with exploratory shifts observed in the gut’s colonization resistance landscape. Ionizing radiation coincided with a notable depletion of key homeostatic commensals and SCFA producers across the cohorts, including *Blautia coccoides, Lactiplantibacillus carotarum, Muribaculum intestinale*, and *Muribaculum gordoncarteri*, alongside mutualistic *Phocaeicola* (*P. dorei, P. sartorii, P. vulgatus*) and Treg-inducing *Clostridium* species (*C. scindens, C. symbiosum*, and sp. C45). This matches the findings from Park et al. (2025), who demonstrated that localized/single-dose irradiation in mice induces severe gut bacteriome dysbiosis, significantly lowers SCFA-producing taxa, triggers intestinal villus damage, and drives pro-inflammatory cytokine surges (IL-6) (10).

The lower representation of *C. scindens* a keystone species that converts primary bile acids into secondary bile acids coincided with an apparent 39% and 41% relative expansion of *Clostridioides difficile* in the cancer and radiation-only groups simultaneously. Because secondary bile acids are the host’s natural defense mechanism to inhibit *C. difficile* germination, radiation-induced dysbiosis effectively shatters colonization resistance.

Pyrogallol administration countered this pathogenic vulnerability by driving the protective *P. distasonis* bloom to outcompete *C. difficile*. Spearman’s rank correlation explicitly connected these dots: pro-inflammatory cytokines (*Il-1α, Il-12*) correlated positively with the pathogenic cluster, likely triggered by lipopolysaccharide (LPS) or pathogen-associated molecular patterns (PAMPs) stimulating toll-like receptor (TLR) signalling across the damaged gut epithelium.

Conversely, the systemic inflammatory cytokine *Il-6* was strongly and negatively associated with the depleted commensals (*M. gordoncarteri, P. dorei, P. vulgatus*, and *P. distasonis*). Because these commensals normally produce SCFAs and indole derivatives (such as indole-3-propionic acid) to preserve the intestinal barrier and stimulate regulatory T cells, their loss removes the "molecular brakes" on host cytokine production.

By enriching *P. distasonis* to a dominant 94% share, pyrogallol co-treatment appears to restore these functional anti-inflammatory brakes, neutralizing pathogen-mediated TLR signalling and downregulating tissue inflammation (*Il-1α, Il-6, Il-12*) during radiotherapy.

### Concluding Remarks and Future Directions

In conclusion, this study demonstrates that pyrogallol functions as a promising multi-target therapeutic adjuvant in oncology. It enhances the efficiency of radiotherapy by driving apoptosis, halting cell-cycle progression, and inhibiting EMT and fibrotic remodelling in host tissues, while concurrently protecting the gastrointestinal tract from radiation-induced inflammatory injury.

While our pooled metagenomic approach mapped an exploratory, representative overview of the microbial landscape in each cohort, it stands as a preliminary investigation that lacks individual intra-group statistical significance. Future research utilizing individual biological replicates is required to capture individual biological variance and validate these taxonomic shifts. Furthermore, integrating untargeted faecal metabolomics will be crucial to decipher how the *Parabacteroides distasonis* bloom alters short-chain fatty acid and bile acid pools to physically drive host tissue repair under carcinogenic conditions.

## Supporting information

gene expression raw data

Primer list

seq metadata mice

species abundance zscores

## Acknowledgement

The authors sincerely thank Dr. Marimuthu Saravanamuthu for his technical expertise and support in executing the localized LINAC radiation protocols for the mice. We also gratefully acknowledge the Indian Knowledge Systems (IKS) division, Ministry of Education, Government of India, for providing the financial support and funding necessary to carry out the mice gut microbiome sequencing and analysis in this study (Sanction order no: 2-23/AICTE/IKS-BGS1/ResearchProject/2022-23/34).

## Ethics approval

All experimental procedures conducted were approved by the Institutional Animal Ethics Committee (IAEC), Sastra Deemed University, Tamil Nadu, India (Approval No. 718/SASTRA/IAEC/RPP).

## Data availability statement

Metagenomic sequencing raw data are publicly available in the NCBI SRA database under BioProject accession PRJNA1504827. All other supporting data, including species abundance and gene expression raw datasets, are available within the article’s Supplementary Material. Any additional underlying raw data will be made available by the corresponding author upon request.

## CRediT authorship contribution statement

**Sayantanee Ray**: Methodology, Formal analysis, Investigation, Visualization, Writing – original draft

**Rubin Nishanth Armstrong**: Methodology, Formal analysis, Investigation, Visualization, Writing – original draft

**Devirpiya Nagarajan**: Conceptualization, Project administration, Validation, Supervision, Writing – review & editing.

**Prakash Shankaran**: Conceptualization, Project administration, Validation, Supervision, Writing – review & editing, Funding acquisition

