## Supplementary material for "Pyrogallol Modulates Abscopal Tumour and Gut Microbial Responses to Localized Irradiation in an Ehrlich Ascites Carcinoma Model": Primer list

| **Genes** | **Forward Primer** | **Reverse Primer** |
| --- | --- | --- |
| Bcl 2 | AGGAGCAGGTGCCTACAAGA | GCATTTTCCCACCACTGTCT |
| Bax | TGCAGAGGATGATTGCTGAC | GATCAGCTCGGGCACTTTAG |
| Caspase 3 | GGGCGTGTTTCTGTTTTGTT | TTGAGGTAGCTGCACTGTGG |
| Caspase 7 | CATATCCACCAGCGCCTTAT | CGCCAGAGGACATGGTTATT |
| CDK 4 | GTGGCTGAAATTGGTGTCGG | TAACAAGGCCACCTCACGAA |
| PCNA | GGAAGCTTAGAGTAGCTCTCATC | GGGAATTCGTGACAGAAAAGACCTC |
| TGF-β | CTCAGACATTCGGGAAGCAGTG | GTATTCCGTCTCCTTGGTTCAGC |
| Col1A1 | AACAACGTCTGCAACTTCGC | CTTCACAAACCGCACACCTG |
| Fibronectin | TCGAGGAGGAAATTCCAATG | ACACACGTGCACCTCATCAT |
| TP53 | GTATTTCACCCTCAAGATCC | TGGGCATCCTTTAACTCTA |
| P21 | CACAGCTCAGTGGACTGGAA | ACCCTAGACCCACAATGCAG |
| N-cadherin | AGGGTGGACGTCATTGTAGC | CTGTTGGGGTCTGTCAGGAT |
| Vimentin | TGAAGGAAGAGATGGCTCGT | TCCAGCAGCTTCCTGTAGGT |
| IL-1α | GCGCTTGAGTCGGCAAAGAAAT | TTCCCGTTGCTTGACGTTGC |
| IL-6 | GACTGATGCTGGTGACAACC | TGGGAGTGGTATCCTCTGTGA |
| IL-12 | CCAGGGAGCATCACGAAGTT | CACTCACTGGTGCTTTGTGC |
| Β-actin | AGCCATGTACGTAGCCATCC | CTCTCAGCTGTGGTGGTGAA |
